# SMCHD1 compacts DNA directly in an ATP-regulated manner

**DOI:** 10.1101/2025.07.08.663435

**Authors:** Joel Ng, Xiaodan Zhao, Xuyao Liu, Aghil Soman, Lars Nordenskiöld, Jie Yan, Shifeng Xue

**Affiliations:** Department of Biological Sciences, National University of Singapore, Singapore; Department of Physics, National University of Singapore, Singapore; School of Biological Sciences, Nanyang Technological University, Singapore; Centre for Bioimaging Sciences, National University of Singapore, Singapore; Mechanobiology Institute, National University of Singapore, Singapore

**Author notes:** Corresponding author: Shifeng Xue.

**Keywords:** SMCHD1, epigenetics, DNA compaction, single-molecule

## Abstract

Structural maintenance of chromosome (SMC) complexes play crucial roles in genome organization by DNA loop extrusion. SMCHD1 is a non-canonical SMC protein important for X-inactivation, imprinting and the silencing of specific autosomal genes. While it is known to be a repressor, little is known about its structure or mechanism of action. Here, we show that the SMCHD1 homodimer is flexible and dynamic in solution. This flexibility is conferred by its yet uncharacterized linker domain, which can alter its length by dynamically switching between compact and extended conformations. Interestingly, we observed that SMCHD1 can directly bridge and compact DNA, independently of other proteins, forming large protein-DNA clusters. SMCHD1 contains a GHKL ATPase domain and SMC hinge domain, however each domain alone is insufficient for DNA compaction. DNA compaction rate decreases when the linker domain is removed. The coiled-coil domain does not affect compaction rate but facilitates interaction with a partner protein LRIF1. Surprisingly, DNA compaction by SMCHD1 does not require ATP and paradoxically, compaction rate is reduced with the addition of ATP. Similarly, SMCHD1 forms clusters with reconstituted nucleosome arrays in the absence of ATP, and the addition of ATP results in a reduction in cluster sizes. Our data provides biophysical and mechanistic insights into the role of SMCHD1 in gene silencing and genome organization.

## INTRODUCTION

Proteins of the Structural Maintenance of Chromosome (SMC) family play important roles in genome organization and gene regulation^1,2^. Structural Maintenance of Chromosomes Hinge-Domain-containing protein 1 (SMCHD1) is a non-canonical SMC protein involved in transcriptional repression, with crucial roles in X chromosome inactivation, genomic imprinting, and silencing of clustered genes^3^. Mutations in SMCHD1 have been attributed to two diseases with currently no known cures: Facioscapulohumeral Muscular Dystrophy (FSHD) and Bosma Arhinia Microphthalmia Syndrome (BAMS)^4–6^. However, the molecular implications of such mutations remain unclear, making it imperative to understand the mechanism of SMCHD1.

While SMCHD1 is known to be required for gene repression, how it achieves this is unclear. Loss of SMCHD1 is generally associated with hypomethylation of target loci, changes in repressive histone marks and 3D DNA conformation^7–12^. However, which effect is directly caused by SMCHD1 and which is secondary is unknown. More recently, SMCHD1 has been characterized as a chromatin insulator and facilitator of long-range interactions^8,9,11^, but how it exerts such functions is also unclear. Therefore, to understand the mechanism of epigenetic repression by SMCHD1, there is a need to determine its direct effects on DNA.

The full-length SMCHD1 protein (2005aa in humans) contains a C-terminus SMC hinge domain and N-terminus GHKL ATPase domain, as well as a lengthy linker region connecting both domains. The SMC hinge domain is found in proteins of the canonical SMC complexes such as cohesin, condensin and SMC5/6; such complexes compact DNA through loop extrusion^13–15^. On the other hand, a GHKL ATPase-containing protein MORC1 was shown in a prior study to compact DNA, albeit through a different mechanism^16^. Based on the presence of these domains, we hypothesize that SMCHD1 has an ability to compact DNA.

Here via single-molecule approaches, we find that human SMCHD1 homodimer is dynamic in solution, conferred by the flexible linker region. When incubated with DNA or nucleosome arrays, SMCHD1 creates dense protein-DNA or protein-nucleosome clusters. Notably, we observe that DNA compaction is independent of ATP. Intriguingly, DNA compaction rate is reduced when the linker region is removed or when ATP is added, and the presence of ATP results in smaller and more dispersed protein-DNA or protein-nucleosome clusters. We further find a role of the coiled-coil domain in interacting with a partner protein LRIF1. Our study provides a first mechanistic insight into how SMCHD1 exerts epigenetic repression.

## RESULTS

### The SMCHD1 homodimer dynamically adopts multiple conformations

While the N-terminus GHKL ATPase domain and C-terminus SMC hinge domain of SMCHD1 have been individually crystallized^17,18^, structural information of the full-length protein has been lacking, other than a negative stain electron microscopy study of mouse SMCHD1^19^. We purified full-length wild-type human SMCHD1 (Fig. 1a) and utilized dry atomic force microscopy (AFM) to visualize its conformation. SMCHD1 purifies as a homodimer as has been observed by past studies^17,20^. Although we observed some SMCHD1 homodimers adopt a closed double rod conformation (Fig. 1a) as similarly reflected by negative stain electron microscopy of mouse SMCHD1^19^, in several AFM scans we observed particles in which the SMCHD1 homodimer appeared in an open conformation (Fig. 1b). Given that the C-terminus hinge domain of SMCHD1 constitutively dimerizes^17,20^, while the GHKL ATPase domain dimerizes only in the presence of ATP^18^, the open conformation likely represents a homodimer connected at the hinge domain. To confirm the appearance of this open-shaped conformation, we measured the lengths of individual particles. The end-to-end length of particles resembling closed double-rod shapes (51.25 nm, n = 73) (Fig. 1c) agrees with the length of the mouse SMCHD1 double-rod homodimer imaged with negative stain electron microscopy (∼40 nm) described previously^19^. The end-to-end length (ATPase domain to ATPase domain) of the open-shaped particles varied based on the angle of hinge domain opening, but the maximum length we observed (∼97nm) is consistent with SMCHD1 adopting an open shaped conformation (Fig. 1c). Our imaging also provided the first indication that the linker region connecting the hinge and ATPase domain is highly flexible and not rigid as previously described for mouse SMCHD1^19^.

**Fig. 1.**
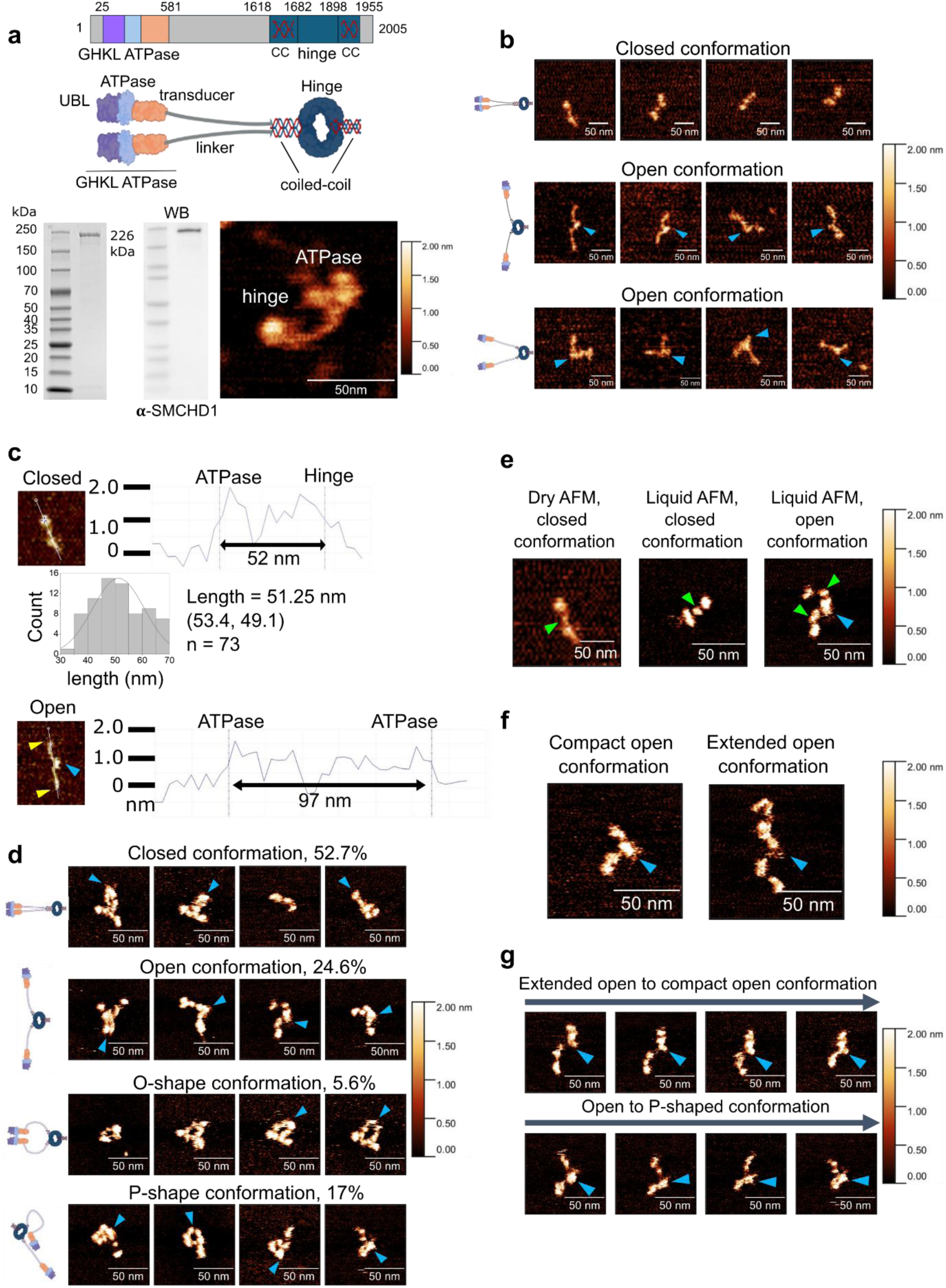
The full length SMCHD1 homodimer adopts various conformations. **a**, *Top*, Domain architecture of full-length SMCHD1 with aa positions corresponding to several constructs used in this study. The SMCHD1 homodimer contains a C-terminus SMC hinge domain, and a N-terminus GHKL ATPase domain, which contains subdomains of a transducer domain, ATPase domain, and UBL (ubiquitin-like domain). *Bottom left*, SDS-PAGE and WB of purified full length SMCHD1. *Bottom right*, representative dry AFM image of a single SMCHD1 homodimer, showing that it adopts a double rod-shape homodimer. **b**, representative images of single SMCHD1 homodimers adopting various conformations in dry AFM, in which we term closed or open conformation. A total of 145 particles were surveyed, with 50.3% and 34.4% adopting closed and open conformations respectively, with the remaining particles not confident of interpretation. **c**, Height analysis via end-to-end section profiles of two SMCHD1 homodimers in either a closed double rod-shaped conformation or open conformation, confirming the open conformation due to increased end-to-end distance. Blue and yellow arrowheads mark the hinge and ATPase domains respectively. Distribution of hinge-to-ATPase distances for the double-rod shape conformation is included, data is presented as means at 95% confidence interval. Histogram is fitted with normal distribution. **d**, Representative images of single SMCHD1 homodimers adopting various conformations in liquid AFM. Two new conformations, P-shape and O-shape (bottom two rows) were identified. Where applicable, blue arrowheads mark the hinge domain. **e**, Dry and liquid AFM images of single SMCHD1 homodimers, with green arrowheads marking the linker central density, and blue arrowhead marking the hinge domain. **f**, *Top*, liquid AFM images of SMCHD1 displaying the linker region in extended or compact conformations. **g**, Successive snapshots from high-speed liquid AFM imaging of SMCHD1, showing that the linker region facilitates SMCHD1 conformational transitions. Where applicable for all AFM images in this figure, blue arrowheads mark the position of the hinge domain.

As dry AFM provides snapshots of protein conformation, we used liquid AFM to assess if the flexibility of the linker domain could allow SMCHD1 to undergo conformational changes in solution. Individual SMCHD1 homodimers were imaged repeatedly under physiological conditions with liquid AFM at a rate of approximately 15 seconds per image (Fig. 1d, Supplementary Fig. 1a, Supplementary Movies 1-3). We observed that SMCHD1 could adopt various conformations (Fig. 1d), and switched between different conformations in real time (Supplementary Movies 1-3). Scanning in liquid also revealed two further conformations which we did not observe in dry AFM, in which here we term as P-shaped and O-shaped conformations (Fig. 1d). The P-shaped conformation depicted one arm of the homodimer being in an extended conformation with the other being in a closed conformation. The O-shaped conformation displayed SMCHD1 as a double-rod closed conformation with the linker domain not entirely associated along its length. We imaged five individual SMCHD1 particles, with at least 100 frames per particle for a total of 1300 frames, and all five individual particles displayed conformational changes over time (Supplementary Fig. 1a, Supplementary Movies 1-3). We quantified the proportion of conformations observed. The closed conformation (52.7%) was the most frequently observed, followed by the open (24.6%), P-shaped (17%) and O-shaped (5.6%).

Liquid AFM also allowed us to visualize further biochemical characteristics of the SMCHD1 linker region. Dry AFM showed the linker to be of low-density and lengthy. Interestingly, in liquid AFM we observed a density clearly visible between the hinge and ATPase domains (Fig. 1e). This linker density was present in the different conformations observed (Supplementary Fig. 1b), but clearest in SMCHD1 particles adopting an open conformation, where the density was visible on both arms of the homodimer. We also observed varying SMCHD1 arm lengths (hinge to ATPase lengths) which suggested that the arm lengths switched between both compact and extended conformations, as revealed by successive snapshots taken during liquid AFM (Fig. 1f, g and Supplementary Fig. 1c). To quantify the typical arm length adopted by SMCHD1, we measured the hinge-to-ATPase lengths in open (n = 321) and P-shape (n = 222) conformations from the liquid AFM images obtained in Figure 1d (Supplementary Fig. 1d). We observed a mean hinge-to-ATPase length of 36.28 ± 0.733 nm (95% confidence interval) and 35.54 ± 1.208 nm (95% confidence interval) for the open and P-shape conformations respectively that was assessed to be not significantly different as expected (Supplementary Fig. 1d).

### SMCHD1 creates dense protein-DNA clusters in an ATP-independent manner

Given the prominent role of SMCHD1 as an epigenetic regulator, we next assessed its potential mechanism by visualizing its interaction with DNA. We incubated 10 nM of SMCHD1 with 1 nM linear 2.6 kb pUC19 DNA substrate for 10 minutes in buffer without ATP, prior to deposition and imaging with dry AFM. Interestingly we observed large clusters formed by SMCHD1-DNA complexes. The clusters were highly dense towards the center while DNA ends could be seen in the periphery protruding from the cluster (Fig. 2a, b). In some cases, multiple DNA ends extended from the cluster, suggesting the cluster can consist of multiple DNA molecules (Fig. 2b). The nature of such center-dense clusters was more evident when a much larger DNA substrate (48.5 kb lambda DNA) was used, where substantial DNA can be clearly seen surrounding a central cluster (Fig. 2c). The volume and maximum z-heights of the DNA clusters were assessed (Fig. 2b). A clear height difference was observed between the clusters (mean 10.2 nm, n = 22), compared to the maximum z-height of SMCHD1 alone (mean 1.17 nm, n = 101) or pUC19 DNA substrate alone (mean 1.10 nm, n = 18). A clear volume difference was also observed, with a mean volume of 117415 nm^3^ for clusters compared to means of 4031 nm^3^ and 666.5 nm^3^ for individual DNA and SMCHD1 particles respectively. Given the low concentrations of SMCHD1 and DNA substrate for our assay, along with our observation that separate imaging of SMCHD1 or DNA alone does not result in any protein aggregation or DNA clustering, we conclude that the dense clusters are induced via the interaction of SMCHD1 with DNA substrates.

**Fig. 2.**
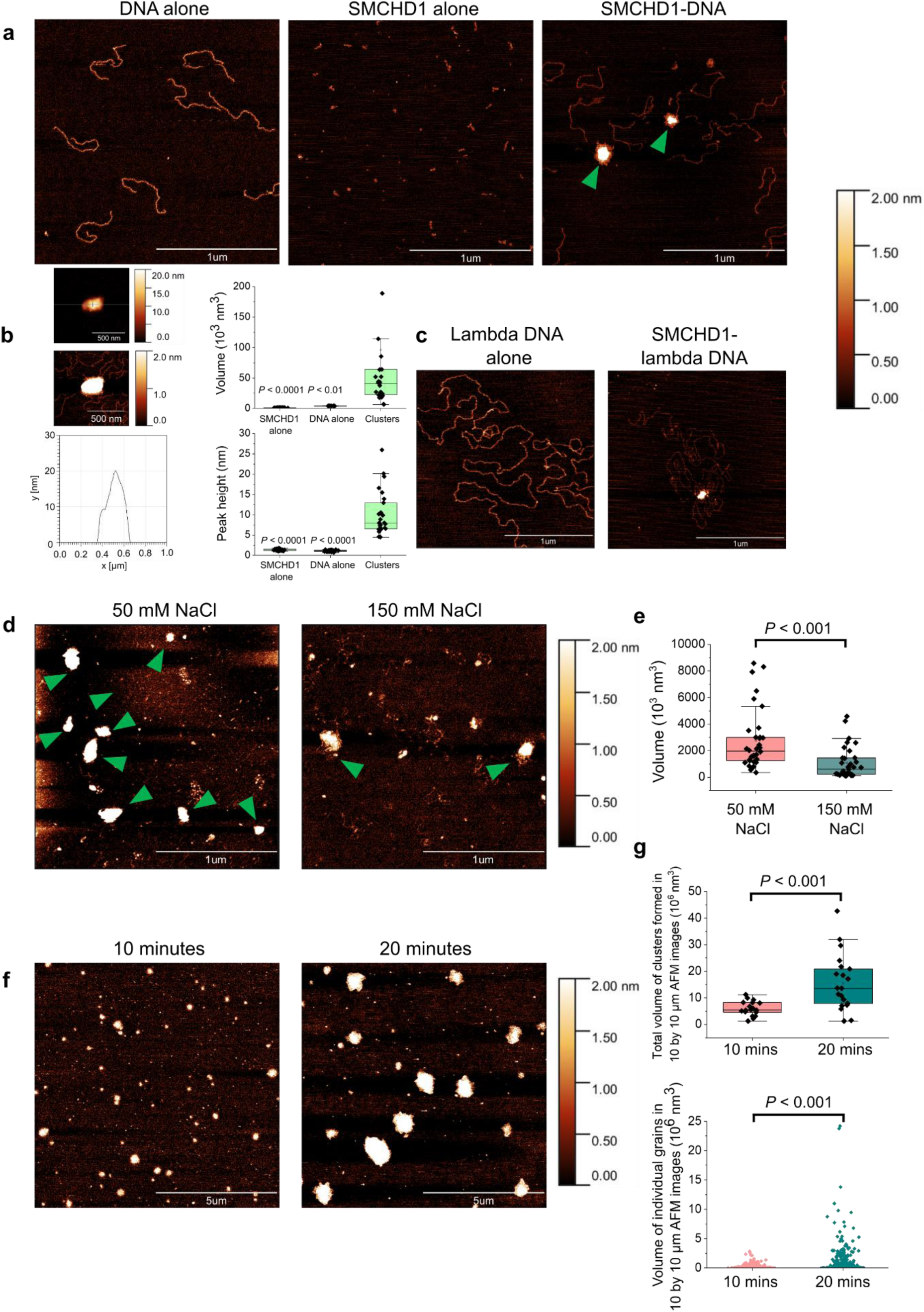
SMCHD1 creates dense protein-DNA clusters. **a**, 2 µm dry AFM images of pUC19 linear DNA alone, SMCHD1 alone, or SMCHD1 incubated with DNA, respectively. Clusters are marked by green arrowheads. **b**, *Left*, horizontal section profile of a single SMCHD1-DNA cluster, with the same AFM image presented at a 20 nm and 2 nm height scale. *Right*, comparison of volume and peak height of individual SMCHD1-DNA clusters, compared to DNA alone or SMCHD1 alone, n = 101, 18, 22 for SMCHD1 alone, DNA alone, and clusters respectively. Volume was measured at a 0.4 nm height threshold. **c**, AFM images of SMCHD1-induced clusters incubated with 48.5 kb lambda DNA substrate. **d**, AFM images of SMCHD1 incubated with pUC19 linear DNA in buffer containing either 50 mM or 150 mM NaCl, deposited on APTES-glutaraldehyde-treated mica. **e**, Volume of clusters assessed from entire 4 by 4 µm AFM images at a 2 nm height threshold, of SMCHD1-DNA clusters in buffer containing either 50 or 150 mM NaCl, n = 31 and 35 respectively, where n indicates individual 4 by 4 µm AFM images, taken from three independent experiments for each condition. **f**, AFM images of SMCHD1 incubated with pUC19 linear DNA in 50 mM NaCl buffer for either 10 or 20 minutes, deposited on APTES-glutaraldehyde-treated mica. **g**, quantification and comparison of total volume of clusters formed in entire 10 by 10 µm images (*top*) or distribution of individual cluster volumes (*bottom*) at a 1 nm height threshold, n = 18 and 21 for 10 and 20 minute conditions respectively, where n indicates individual 10 by 10 µm AFM images from two independent experiments. For boxplots in b, e and g, the middle quartile marks the median, boxes extend to the quartiles, and the whiskers depict the range of the data within 1.5 times of the interquartile range of the median; two tailed t-test was used for statistical analysis. Source data is provided as a source data file.

To investigate the effect of ionic strength on SMCHD1-DNA clusters, we set up similar SMCHD1-DNA incubations in buffer containing either 50 or 150 mM NaCl and deposited the reactions on APTES-glutaraldehyde functionalized mica; such a surface-modified mica allows for DNA-protein complexes to be deposited under a range of buffer conditions^21,22^. We observed visible decrease in intensity of clusters in 150 mM NaCl compared to 50 mM NaCl (Fig. 2d). We quantified the total volume of clusters formed per 4 by 4 µm AFM images and observed a significant difference in total cluster volumes, with means of 1.03 x 10^6^ nm^3^ and 2.88 x 10^6^ nm^3^ for 150 mM and 50 mM NaCl respectively (Fig. 2e), whilst depositing and imaging SMCHD1 alone or DNA alone in either ionic condition did not result in aberrant aggregation or clustering (Supplementary Fig. 2a). Our results suggest that formation of clusters by SMCHD1 is regulated by ionic strength. Nevertheless, our observation of cluster formation at physiological salt concentration of 150 mM NaCl suggests that cluster formation is relevant *in vivo*. Using the similar APTES-glutaraldehyde deposition method, we also observed that the volume of clusters visibly increased in a time-dependent manner (10 vs 20 minutes) (Fig. 2f, g).

Lastly, to completely rule out any potential effects induced by counterion interference, we incubated SMCHD1 and DNA in counterion-free buffer (20 mM Tris-Cl pH 8.0, 50 or 150 mM NaCl, 0.5 mM TCEP) and deposited on freshly cleaved unmodified mica. The absence of the counterion would prevent any free DNA from adhering to the negatively charged mica surface but still allowed protein-containing mixtures to be visualized. As expected, we did not observe any free unbound DNA but still noticed dense DNA clusters at both 50 mM and 150 mM NaCl conditions (Supplementary Fig. 2b). Likewise, deposition and imaging of SMCHD1 alone resulted only in monodispersed particles without any clustering (Supplementary Fig. 2b). These observations confirm that SMCHD1 is monodisperse when imaged alone, while forming clusters in the presence of DNA.

### SMCHD1 creates clusters by compacting and bridging DNA molecules

To understand the mechanism of SMCHD1-DNA cluster formation, we sought to capture intermediates of cluster formation. The concentration of SMCHD1 was reduced to 5 nM, and the SMCHD1-DNA incubation time reduced from 10 minutes to 1 minute, to allow visualization of single SMCHD1 particles on DNA. The C-terminus hinge domain of SMCHD1 has been shown to bind DNA^23^. Much less is known regarding the DNA-binding characteristics of the N-terminus GHKL ATPase domain, although the GHKL ATPase domain of MORC3 can bind DNA^24,25^. We obtained scans of single SMCHD1 homodimers bound to DNA substrates (Fig. 3a). Interestingly we observed the ATPase domain in contact with DNA (Fig. 3a, left), thus suggesting that the full-length protein could act as a bridge to bring adjacent segments of DNA into close proximity. Clusters could be localized towards DNA ends or in the middle of the DNA molecules; SMCHD1 binding could also bring about DNA bending and connection of adjacent regions of single DNA molecules (Fig. 3b).

**Fig. 3.**
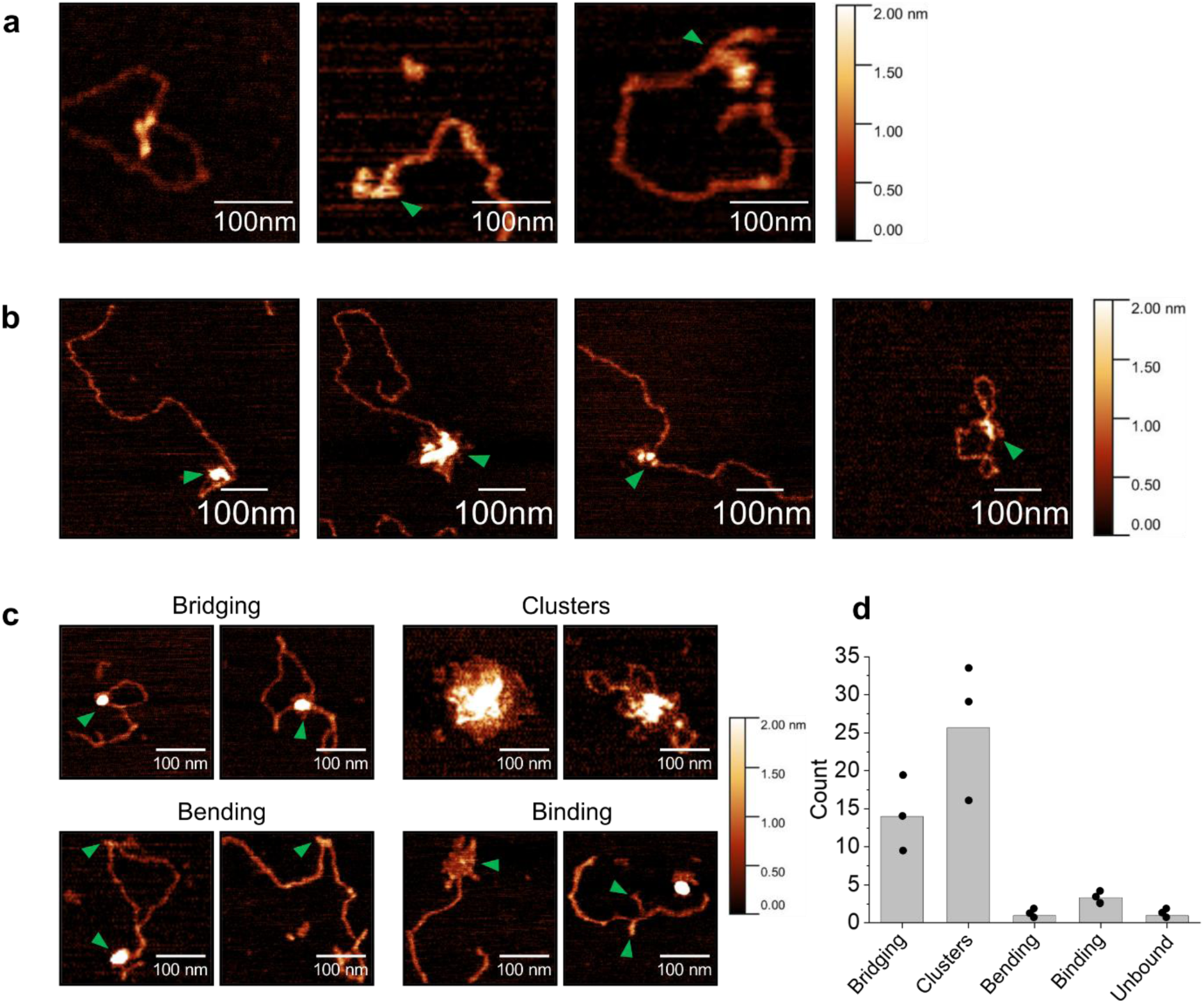
SMCHD1 creates clusters by compacting and bridging DNA. **a**, Dry AFM images of a single SMCHD1 homodimer bound to DNA. **b**, AFM images of DNA substrates with SMCHD1-induced clustering localized towards the DNA ends, and AFM images showing that SMCHD1 is able to facilitate DNA bending and bridging. **c**, representative AFM images of various SMCHD1 interaction events, with green arrowheads indicate bridging, bending or binding events. **d**, quantification of SMCHD1 interaction events from (c) indicating DNA bridging, DNA clustering, DNA bending, DNA binding (without bridging or clustering), and finally unbound DNA. Bars indicate mean with individual data points taken from three individual experiments (n = 35, 44, 56) for a total of 135 individual DNA molecules/clusters. The 2nm z-scale bar applies to all images in this Figure.

To quantify this bridging phenomenon, we assessed individual DNA molecules and classified them into whether we observed (1) bridging, (2) cluster formation, (3), bound SMCHD1 but no bridging or clustering, or (4) unbound DNA (Fig. 3c). We observed bridging and clustering events at 29.2% and 54.7% respectively (Fig. 3d). Given that SMCHD1 was incubated with DNA in the absence of ATP, these DNA bridging and clustering events we observe are unlikely to be ATP-dependent loop extrusion. Instead, we propose that SMCHD1 binds to DNA through different domains, and joins adjacent DNA regions together.

### SMCHD1 hinge and ATPase domains alone do not induce clusters

SMCHD1 contains a C-terminus hinge domain and N-terminus ATPase domain. We investigated the individual contributions of these domains alone towards the observed SMCHD1-DNA cluster formation. We purified the hinge domain (aa1682 to 1898), ATPase domain (aa25 to 581) and extended ATPase domain (aa25 to 702) individually (Fig. 4a) and subjected them to a similar AFM protocol as the full-length protein in figure 2a. Although DNA binding was observed for the hinge domain as expected, even an excess of protein deposited (50 nM) failed to induce cluster formation (Fig. 4b). No prior study has investigated the DNA-binding properties of the ATPase domain. As we observed the ATPase domain of full-length SMCHD1 to be in contact with DNA (Fig. 3a, left), we performed an electrophoretic mobility shift assay (EMSA) to confirm its DNA binding ability. Surprisingly the ATPase domain also resulted in gel shift of DNA (Fig. 4c). In our initial AFM analysis using 10 nM of the ATPase domain, we observed binding of the ATPase domain to DNA, with binding events coinciding with the bending of DNA but no compaction (Fig. 4d). Interestingly, at 50 nM, the ATPase domain could create localized minor clusters at individual DNA molecules, along with the ability to join multiple DNA molecules together (Fig. 4e). We observed a similar phenomenon with the extended ATPase domain (aa25-702) (Fig. 4f). Nevertheless, even at excess concentrations for the hinge and ATPase domains individually at 50 nM, we did not observe any dense cluster formation as we did observe with full-length SMCHD1 (10 nM used). This shows that neither domain is sufficient by itself to compact DNA into dense protein-DNA clusters.

**Fig. 4.**
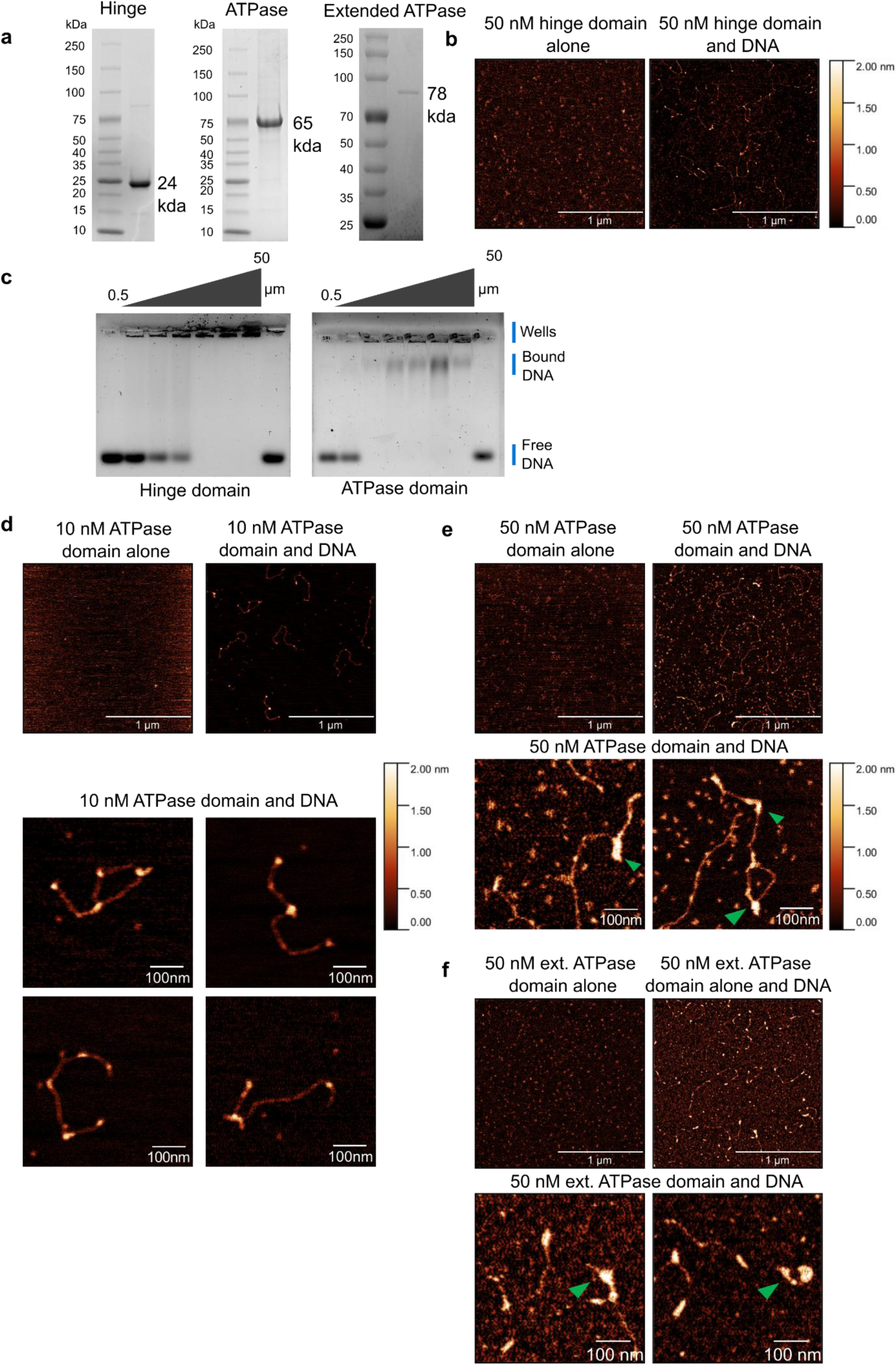
SMCHD1 hinge and ATPase domains bind DNA but do not induce clusters. **a**, SDS-PAGE of purified SMCHD1 hinge domain (aa1682-1898), ATPase domain (aa25-aa581) and extended ATPase domain (aa25-702). **b**, Dry AFM images of either 50 nM hinge domain alone or incubated with DNA. Binding of the hinge domain to DNA is observed without cluster formation. **c**, EMSA analysis for purified hinge and ATPase domains. Concentration of protein used: 0.5, 10, 20, 30, 40, 50 µM. The first and last lanes of the gel are loaded with free DNA. **d**, AFM images of 10 nM of ATPase domain (aa25-581) alone or with DNA. **e**, AFM images of 50 nM of SMCHD1 ATPase domain (aa25-581) alone or with DNA. **f**, AFM images of 50 nM of extended ATPase domain (aa25-702) alone or with DNA. Where observable, key points of contact of protein with DNA are indicated with green arrowheads.

### The coiled-coil and linker domains of SMCHD1 are not required for compaction

Given the observation that SMCHD1 adopts multiple conformations, we assessed if the inherent flexibility of SMCHD1 is linked to its function in creating protein-DNA clusters. The full-length SMCHD1 protein contains minor coiled-coil domains on both sides of the C-terminus hinge domain. Coiled-coil domains are highly prevalent in the canonical SMC core proteins (SMC1-6), serving as a linker region which connect the hinge and ATPase heads, and have been shown to confer flexibility^26–29^. Mutation of lysine residues within the coiled-coils have also been shown to abrogate inter-coiled-coil association^28^. In SMCHD1, the combined size of the coiled-coil domains (∼22 kDa) is minor relative to the canonical SMC proteins. Given our observation of inherent flexibility in the full-length SMCHD1 protein, we first investigated if the coiled-coil domains in SMCHD1 had any effect on the flexibility and functionality of the full-length protein.

We purified a truncated SMCHD1 (from here on denoted SMCHD1^△CC^) in which we removed both coiled-coil domains (removal of aa1618-1682 and aa1900-1955) and assessed its conformation with AFM. Using similar liquid AFM imaging as for wild-type SMCHD1 (from here on denoted SMCHD1^WT^), we observed that SMCHD1^△CC^ was still able to undergo dynamic changes in conformations (Fig. 5a and Supplementary Fig. 3). Furthermore, we also observed that SMCHD1^△CC^ could still compact DNA and form dense DNA clusters (Fig. 5b, c). A second albeit less prevalent function of coiled-coil domains is protein-protein interaction. For instance, the coiled-coil region of SMC5 was shown to be required for interaction with the E3 ligase Mms21^30^. As such we assessed if the coiled-coil domain of SMCHD1 exhibited a similar function with a known interactor LRIF1^31^. LRIF1 is a validated direct interactor of SMCHD1 and has been partially attributed to SMCHD1 localization to the inactive X chromosome^19,31^. A nonsense mutation in LRIF1 has also been implicated in FSHD^32^. In 293T cells, we observed that immunoprecipitation of LRIF1 was significantly reduced with SMCHD1^△CC^ as compared to SMCHD1^WT^ (Fig. 5d), suggesting that the coiled-coil domain of SMCHD1 is required for protein-protein interaction rather than conferring flexibility or DNA compaction.

**Fig. 5.**
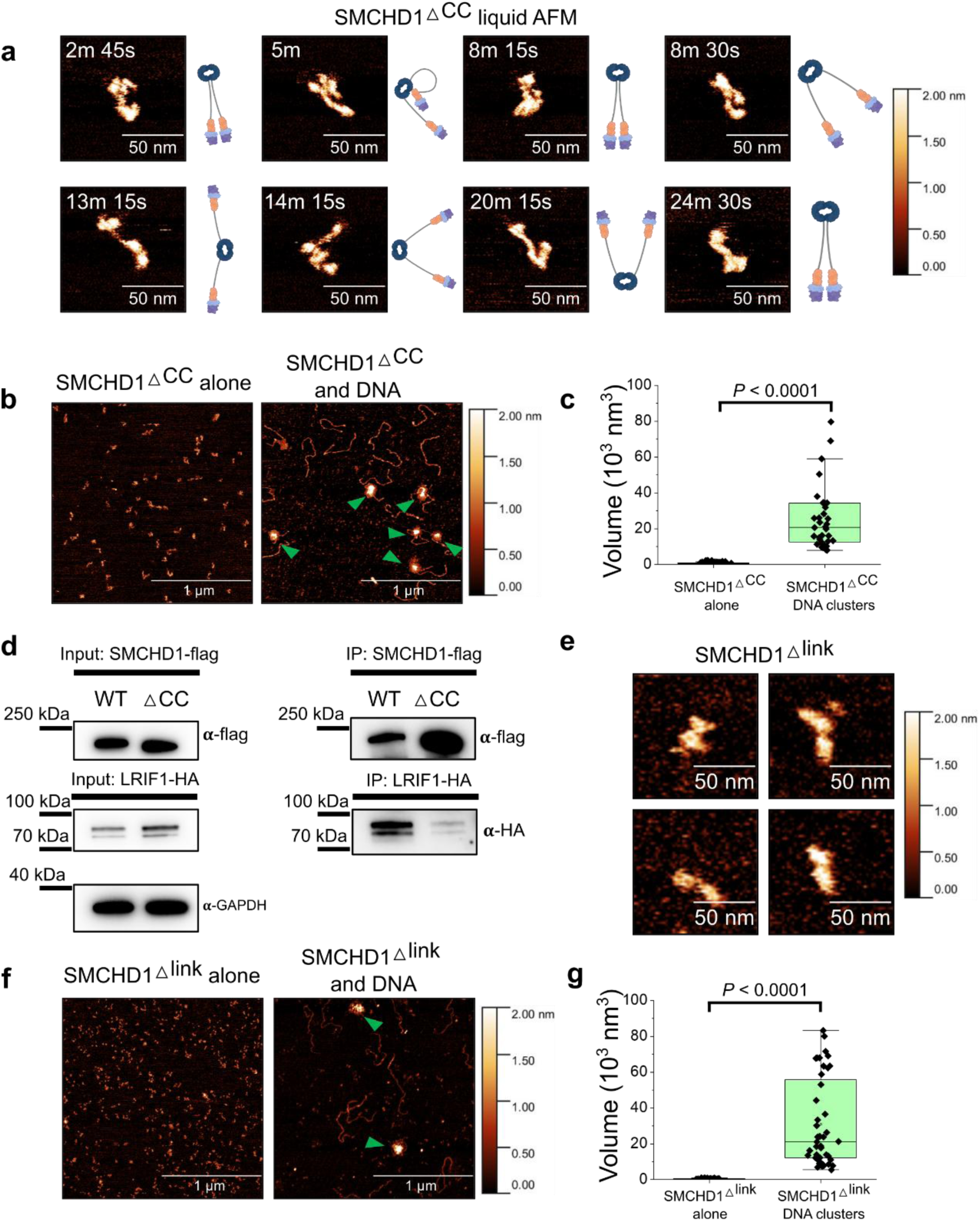
Evaluation of the coiled-coil and linker domains in SMCHD1 function. **a**, Liquid AFM images of an individual SMCHD1^△CC^ homodimer adopting various conformations. Timepoints indicate the snapshots taken during the imaging process. Two molecules were investigated for SMCHD1^△CC^ totaling 201 frames, with closed (48.3%), open (23.9%), P-shape (9.45%) conformations observed, 18.4% of frames were not confident for interpretation. **b**, Dry AFM images of SMCHD1^△CC^ alone or DNA clusters formed by SMCHD1^△CC^, with clusters marked by green arrowheads. **c**, comparison of volumes of SMCHD1^△CC^ particles alone (n = 171) compared to individual SMCHD1^△CC^-DNA clusters (n = 40) at a 0.4 nm height threshold. **d**, Western Blot analysis of flag-IP in 293T cells demonstrating co-immunoprecipitation of LRIF1-HA with SMCHD1-flag. Three independent experiments were conducted **e**, Liquid AFM images of SMCHD1^△link^. **f**, Dry AFM images of SMCHD1^△link^ alone or DNA clusters formed by SMCHD1^△link^, with clusters marked by green arrowheads. **g**, comparison of volumes of individual SMCHD1^△link^-DNA clusters (n = 47) compared to individual SMCHD1^△link^ particles alone (n = 59) at a 0.4 nm height threshold. For c and g, two tailed t-test was used for statistical comparison. Source data is provided as a source data file.

To determine the function of the lengthy linker in SMCHD1, we purified another truncated version of SMCHD1 by removing the entire linker (removal of aa703-1615) but retaining the coiled-coil domains (from here on denoted SMCHD1^△link^). As expected, as we had removed ∼101 kDa the resulting truncated protein no longer resembled a rod-like particle. Nevertheless, we were able to visualize SMCHD1^△link^ as two globular domains side-by-side, corresponding to the hinge and ATPase domains (Fig. 5e). SMCHD1^△link^ could still create dense protein-DNA clusters as assessed with AFM, indicating that the linker is also dispensable for cluster formation under these conditions (Fig. 5f, g).

### SMCHD1 compacts DNA molecules against low physical forces

Given the limitations of AFM in assessing DNA compaction in real-time, we utilized magnetic tweezers to gain a more dynamic view of DNA compaction brought about by SMCHD1. Linear 7.5 kb lambda DNA substrates were tethered onto glass slides of flow channels and stretched with a vertical magnetic tweezers setup (Fig. 6a). Experiments were conducted by stretching DNA at a high force (8 pN) before injecting 10 nM SMCHD1, followed by observing compaction at a constant low force (0.5 pN). This force was chosen based on prior loop extrusion studies of canonical SMC complexes, for instance 0.3-0.6 pN shown for cohesin^14^.

**Fig. 6.**
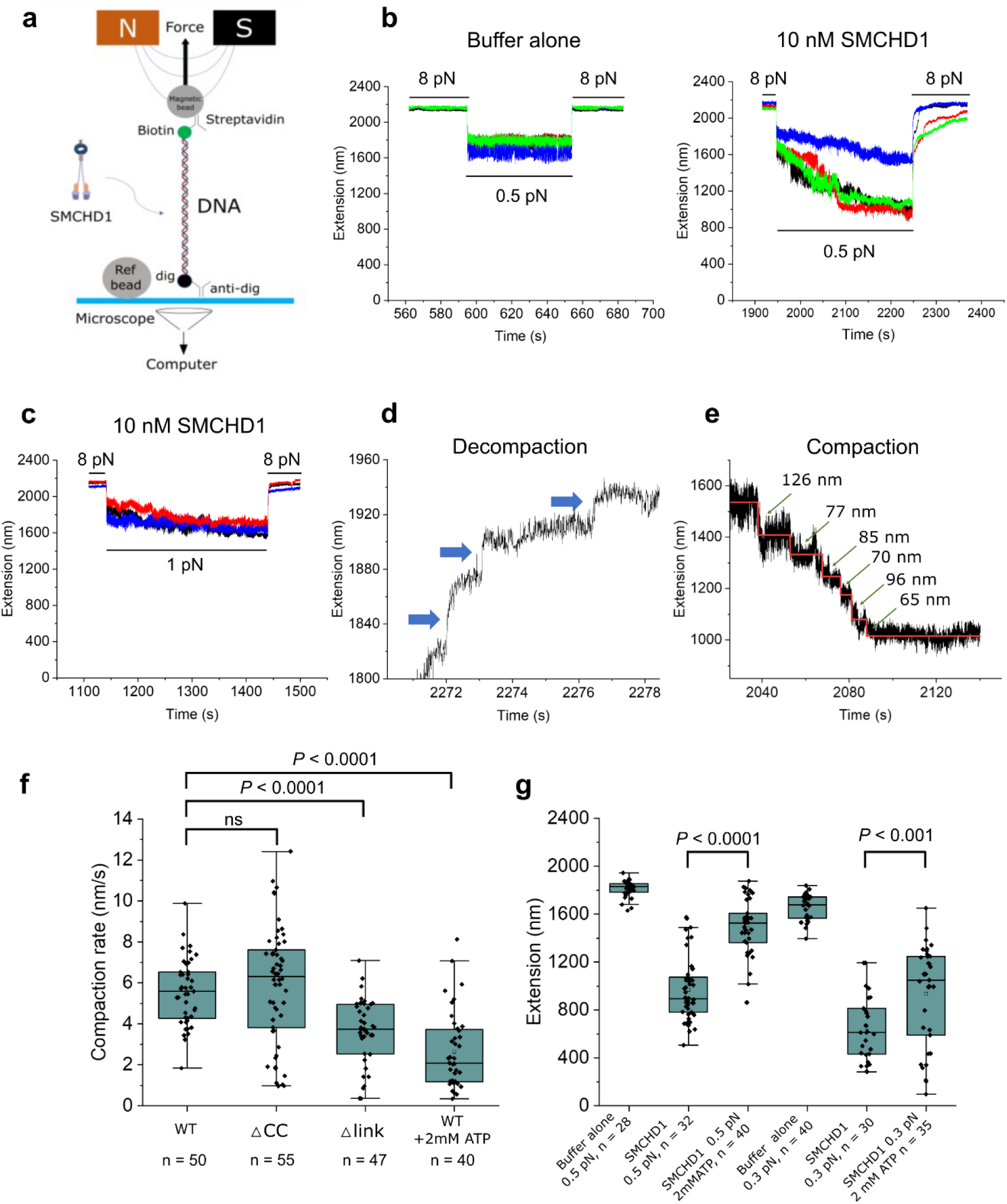
SMCHD1 compacts DNA at low forces in magnetic tweezers assays. **a**, Experimental configuration of vertical magnetic tweezers set up. DNA substrates are tethered via anti-digoxigenin to one end (glass base) and biotin interaction to streptavidin magnetic beads. An overhead magnetic applies force to the magnetic beads to stretch DNA. **b**, *Left*, sample control (buffer alone) extension of DNA substrate to demonstrate the change in DNA length at 8 pN and 0.5 pN of force. *Right*, compaction traces of 10 nM SMCHD1 at 0.5 pN. Each color trace represents an individual DNA tether. **c**, Similar compaction trace at 1 pN of force. Each color trace represents an individual DNA tether. **d**, Representative decompaction trace from **b** denoting the discrete steps (marked by blue arrows) observed as the DNA extends to its original length under high force. **e**, Representative compaction trace of 10 nM SMCHD1-mediated DNA compaction at 0.5 pN, with the red line indicating the fit by Autostepfinder^33^ and arrows indicating the step sizes. **f**, Box-plot comparison of DNA compaction rates between SMCHD1^WT^, SMCHD1^△CC^, SMCHD1^△link^, and finally SMCHD1^WT^ with 2 mM ATP. The middle quartile marks the median, boxes extend to the quartiles, and the whiskers depict the range of the data within 1.5 times of the interquartile range of the median. Statistical analysis was performed by one-way ANOVA with Dunnett’s test for multiple comparisons. **g**, Box plot comparison of DNA extension lengths between SMCHD1 with or without ATP at either 0.5 or 0.3 pN of force, recorded at the end of the 81s duration of compaction. Control extension traces (buffer alone) are included. The median, quartiles, and data range are as described in **f**. Statistical analysis was performed using unpaired two-tailed t-test. Data in **f-g** are indicated (*n*) events collected from at least three independent measurements for each condition, where *n* indicates individual DNA tethers. Source data is provided as a source data file.

We observed that SMCHD1 could compact DNA best at forces of 0.5 pN and below (Fig. 6b), albeit with marginal compaction at 1 pN (Fig. 6c). This compaction was reversible as a large force (8 pN) could restore DNA length. However, force-induced decompaction occurred gradually and did not immediately restore DNA to its original length. On closer examination, we observed discrete decompaction steps ranging between 50 to 100 nM of DNA length (Fig. 6d). These observations suggest that SMCHD1 compacts DNA by bridging together adjacent DNA segments to form multiple loops, such that during force-induced decompaction, these loops are disrupted to account for the distinct decompaction steps. To obtain a more robust analysis of such loop formation, we fitted compaction traces with an automated bias-free algorithm Autostepfinder^33^. At forces of 0.5 pN and 0.3 pN, we observed fitted compaction steps with a median of 45 ± 20.3 and 90 ± 42.6 nm (median absolute deviation (MAD)) respectively, which suggests that SMCHD1 compacts DNA by bridging adjacent DNA segments (Fig. 6e, Supplementary Fig. 4).

As magnetic tweezer system provides more dynamic measurements than AFM, we conducted similar experiments with SMCHD1^△CC^ and SMCHD1^△link^ and assessed DNA compaction rate compared to SMCHD1^WT^ over an experimental duration of 81 seconds. Surprisingly, while removal of the coiled-coil domains in SMCHD1^△CC^ did not significantly affect compaction, removal of the linker domain in SMCHD1^△link^ significantly reduced compaction rate (Fig. 6f). Interestingly, the addition of ATP in magnetic tweezers assays also significantly reduced the DNA compaction rate of SMCHD1 when conducted at a force of 0.5 pN (Fig. 6f). To further validate this observation, we repeated the comparison at a force of 0.3 pN instead of 0.5 pN and recorded the final DNA extension length at the end of the compaction duration of 81s, with similar observations (Fig. 6g). Taken together, while SMCHD1 compacts DNA in all conditions tested, compaction rate is reduced with the removal of the linker domain or with the addition of ATP.

### SMCHD1 forms clusters with nucleosomal DNA

While loop extrusion assays with canonical SMC complexes have typically utilized naked DNA, several studies have also demonstrated that the canonical SMC complexes can compact nucleosome-bound DNA^14,34,35^. As SMCHD1 has been characterized as a mediator of chromatin interactions^8,9,11^, we investigated whether SMCHD1 can compact nucleosomal DNA.

We reconstituted nucleosome arrays, using a 15-repeat Widom 601 sequence with 50 bp linker DNA (denoted 15-197-601) (Fig. 7a and 7b). Using AFM, we observed that SMCHD1 formed clusters with reconstituted nucleosome arrays that increased in size in an SMCHD1 concentration-dependent manner (20, 40 and 80 nM) (Fig. 7c), with no clustering observed in SMCHD1-alone controls (Supplementary Fig. 5a). Interestingly with the addition of ATP, we observed that SMCHD1-nucleosome clusters were smaller and more dispersed (Fig. 7d), with no differences in nucleosome array-alone controls (Supplementary Fig. 5b). The height and volume of individual clusters also increased in a SMCHD1 concentration-dependent manner (Supplementary Fig. 5c). The crystal structure of the catalytically inactive SMCHD1 ATPase domain alone showed that the ATPase domain dimerizes upon ATP binding^18^. To verify that the increased number and highly dispersed nature of clusters was not due to aberrant SMCHD1 clustering or dimerization in the presence of ATP, we assessed the volume of individual SMCHD1 particles with 20 nM concentration of SMCHD1 alone, with and without ATP using liquid AFM (Supplementary Fig. 5d). There was no indication of a doubling in the mean volume of individual SMCHD1 particles in the presence of ATP. This suggests that the presence of ATP did not result in oligomerization of SMCHD1 and that the identity of the smaller dispersed clusters we observed was not due to aberrant SMCHD1 multimers. Furthermore, to rule out any negative charge effect of ATP, we used GTP instead of ATP. This resulted in the formation of clusters which more resembled the ATP-null condition rather than the ATP condition (Supplementary Fig. 5e). This demonstrates that the dispersed clusters observed with ATP is a SMCHD1-ATP-specific phenomenon.

**Figure 7.**
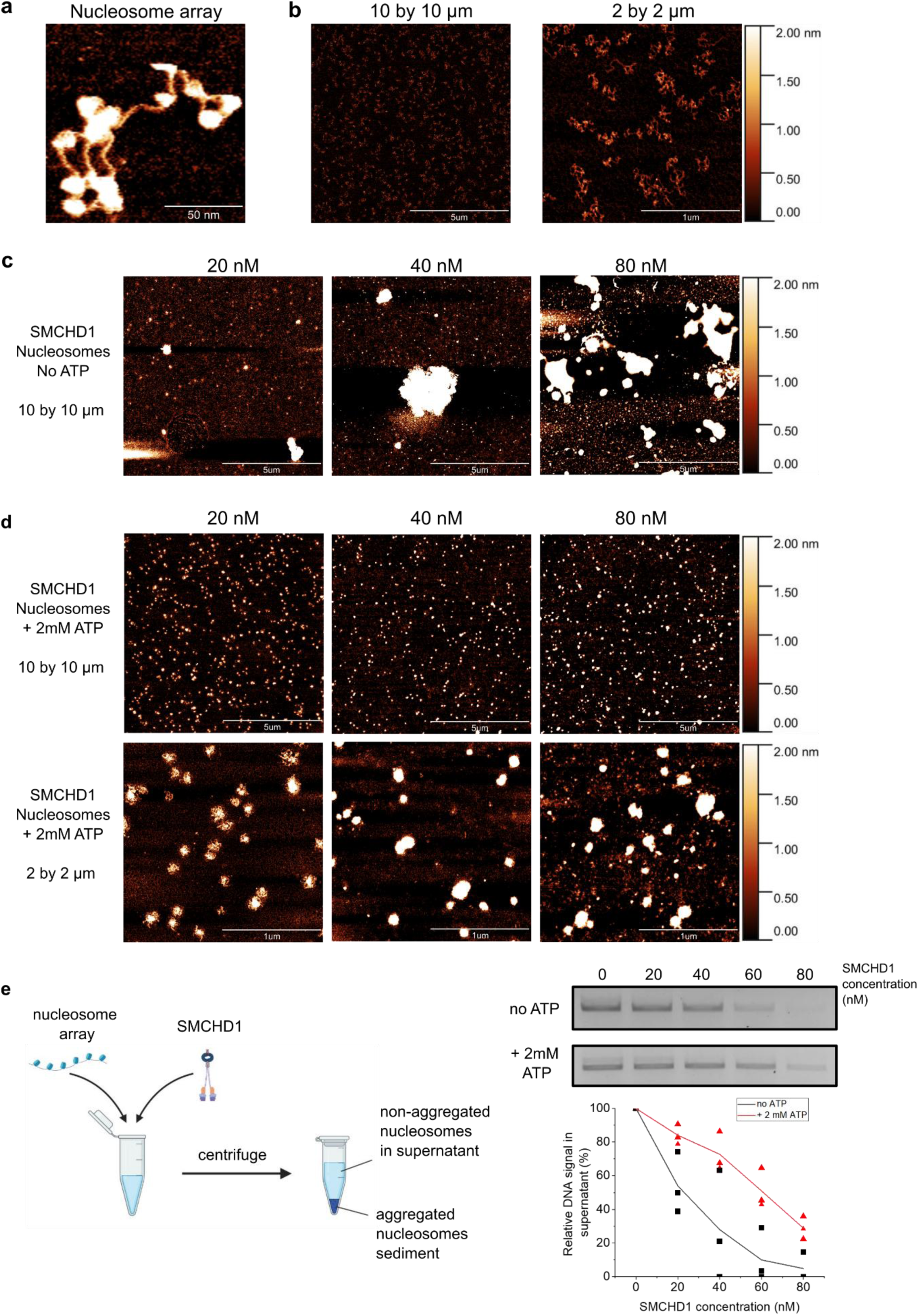
SMCHD1 forms clusters with reconstituted nucleosome arrays. **a**, 2 by 2 µm liquid AFM image of a single nucleosome array. **b**, 10 by 10 µm and 2 by 2 µm AFM images of nucleosome arrays alone deposited on poly-L-lysine treated mica. **c**, 10 by 10 µm AFM images of SMCHD1-nucleosomes at indicated SMCHD1 concentrations (20, 40, 80 nM SMCHD1) in the absence of ATP. **d**, 10 by 10 µm (top row) and 2 by 2 µm (bottom row) AFM images of SMCHD1-nucleosomes at indicated SMCHD1 concentrations (20, 40, 80 nM SMCHD1) in the presence of ATP. **e**, Polynucleosome association assay. *Left*, schematic of the polynucleosome association assay. Nucleosome arrays are incubated with SMCHD1 at indicated concentrations (20 to 80 nM) in the absence or presence of ATP, and then isolated with centrifugation. The supernatant containing non-aggregated nucleosomes can be assessed with gel electrophoresis. *Right*, agarose gel electrophoresis and quantification of supernatant containing non-aggregated nucleosomes, at indicated SMCHD1 concentrations in the absence or presence of ATP. DNA band intensities were assessed with ImageJ and plotted relative to the band corresponding to 0 nM SMCHD1 added. Three independent experiments were conducted, each data point represents a band intensity, and the line represents the average of the band intensities.

To validate the observed ATP-dependent effects, we also utilized a modified polynucleosome association assay^36^, which assesses the amount of nucleosomes remaining in solution following incubation with SMCHD1 and subsequent sedimentation (Fig. 7e). This bulk assay complements our AFM observations by providing a sample-wide assessment. We observed that with increasing SMCHD1 concentrations (20 to 80 nM), the amount of nucleosomes in solution progressively decreased, attributed to the increased sedimentation of SMCHD1-nucleosome clusters. In the presence of ATP, more nucleosomes remained in solution, confirming a decrease in the degree of clustering and sedimentation (Fig. 7e). In combination with our observations in AFM, our results demonstrate that SMCHD1 forms clusters with nucleosomes, and addition of ATP results in an increased number of dispersed clusters of reduced volume.

### Effect of radicicol on nucleosomal compaction

To further validate the effects of ATP-dependent action of SMCHD1 on nucleosomal compaction, we conducted similar experiments using 80 nM SMCHD1 together with the GHKL-ATPase specific inhibitor radicicol. An ATPase assay revealed a mean 48% reduction in ATP-hydrolysis rate compared to the radicicol-null condition (Fig. 8a). In SMCHD1 nucleosome AFM experiments with ATP and radicicol, clusters formed an intermediate size between the ATP-null and ATP condition, with individual clusters smaller compared to the ATP-null condition, but larger than the ATP condition (Fig. 8b compared to Fig. 7c, d). To quantify between all conditions, we marked grains using a 2 nm height threshold and assessed grain distributions from multiple 10 by 10 µm AFM images from ATP-null (n = 29 images), ATP-radicicol (n = 33 images), and ATP (n = 35 images) conditions (Fig. 8c). We observed significant differences in grain distribution volumes between ATP-null and ATP-radicicol conditions, as well as between ATP-radicicol and ATP conditions (Fig. 8c), confirming an intermediate volume formed in the ATP-radicicol condition between the ATP-null and ATP condition.

**Figure 8.**
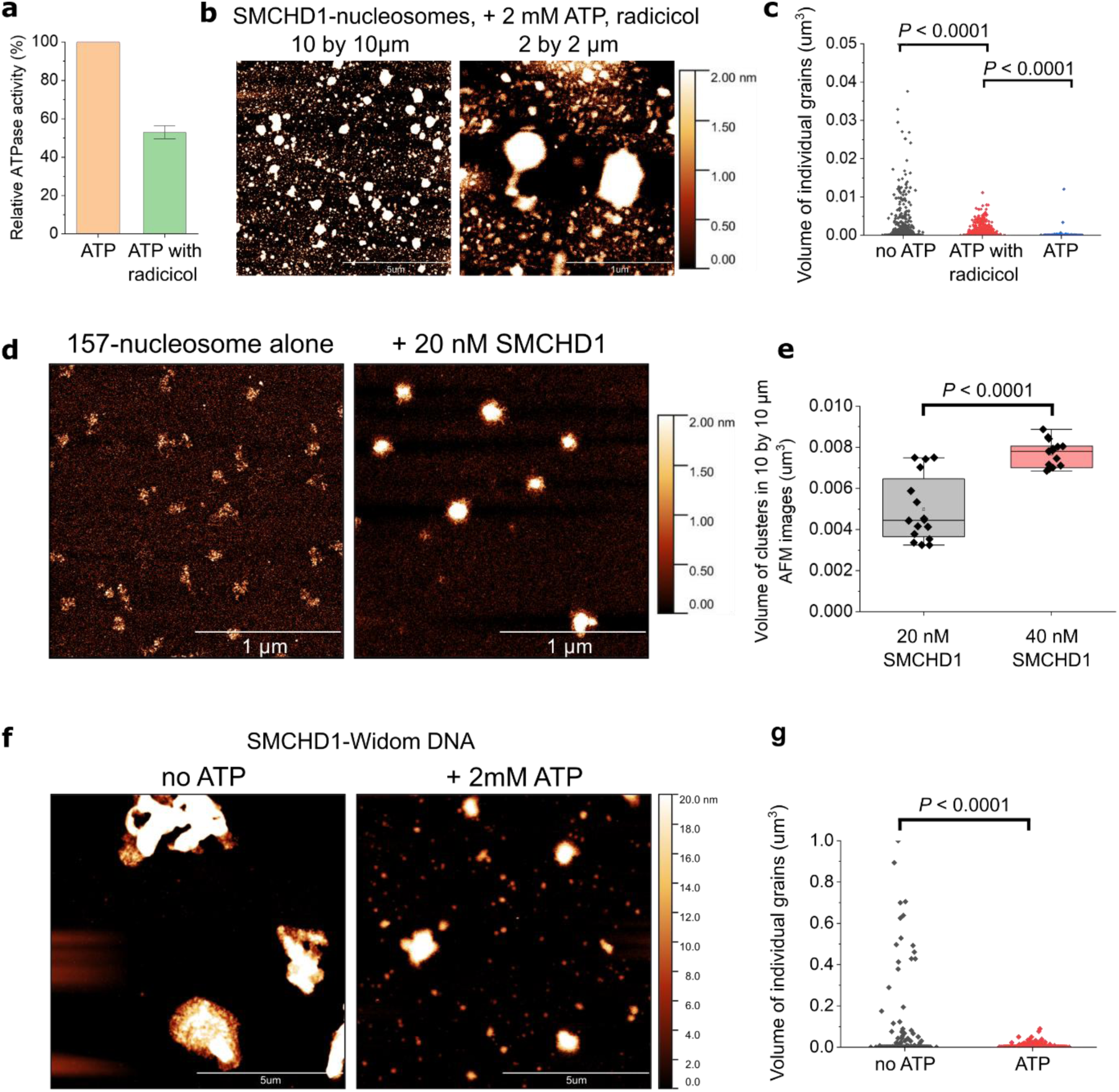
Effects of radicicol and linker DNA on SMCHD1 nucleosome clusters. **a**, ATPase assay conducted as indicated, demonstrating a decrease in ATPase hydrolysis rate in the presence of 100 µM radicicol. Bars represent mean with standard deviation. **b**, 10 by 10 and 2 by 2 µm images of 80 nM SMCHD1 in the presence of ATP and radicicol. **c**, comparison of grain distributions generated by gwyddion at a 2 nm height threshold for 10 by 10 µm AFM images, comparing between ATP-null (n = 29 images), ATP-radicicol (n = 33 images), ATP (n = 35 images) from three independent experiments for each condition. For better visualization and comparison the range of graph in the y axis was limited up to 0.05 um^3^ (see source data). **d**, 2 by 2 µm AFM images of SMCHD1 nucleosome clusters using a 20-157-601 nucleosome array that has significantly shorter linker DNA length of 10bp. **e**, boxplot comparisons of the total volume of clusters between 20-157-601 clusters formed between 20 or 40 nM SMCHD1, n = 16 for both conditions across two independent experiments, where n is an individual 10 by 10 µm AFM image. **f**, 10 by 10 µm AFM images of 80 nM SMCHD1 incubated with Widom 601 naked DNA that was used for reconstitution of nucleosomes. **g**, comparison of grain distributions generated by gwyddion formed in multiple 10 by 10 µm AFM images for ATP-null (n = 26 images) and ATP (n = 35 images) conditions across three independent experiments for each condition. For better visualization and comparison the range of graph in the y axis was limited up to 1 um^3^ (see source data). The 2 nm z-height scale bar applies to all AFM images in this figure, with the exception of **f** in which a 20 nm z-height scale bar is used. Two tailed t-test was used for statistical analysis (**c**, **e**, **g**). For the boxplots in **e**, the middle quartile marks the median, boxes extend to the quartiles, and the whiskers depict the range of the data within 1.5 times of the interquartile range of the median. Source data is provided as a source data file.

### Cluster formation and regulation by ATP does not depend on linker length or nucleosomes

As a next measure of validating nucleosome compaction by SMCHD1, we utilized nucleosome arrays with significantly shorter linker DNA (10 bp) and observed similar cluster formation in the presence of SMCHD1 and ATP (Figure 8d). We quantified the total volume of clusters from multiple 10 by 10 µm AFM images and observed an increase in volume from 20 to 40 nM SMCHD1 (Figure 8e). Finally, we tested if the observed effect of ATP-dependent disperse clusters was consistent on naked DNA. We conducted a similar experiment using 80 nM of SMCHD1 incubated with the Widom-601 naked DNA used for nucleosome reconstitution, in the presence and absence of ATP. Similar to experiments with nucleosome arrays, we observed a similar ATP-dependent phenomenon, in which ATP led to visibly disperse clusters compared to the ATP-null condition (Fig. 8f). We quantified the volume of all clusters formed in multiple 10 by 10 µm images and observed a significance decrease in the volume of clusters formed (Fig. 8g). Our results thus demonstrate that SMCHD1-induced cluster formation is a reproducible phenomenon irrespective of naked DNA or chromatin that may differ in linker DNA length, with the cluster sizes regulated by ATP.

## DISCUSSION

Using a combination of single-molecule and biochemical experiments, we provide key insights into a possible epigenetic mechanism of SMCHD1. To date, the majority of studies assessing SMCHD1 function have relied on genomic studies such as chromosome conformation capture (Hi-C) and gene expression, with a consensus that SMCHD1 is involved in chromatin remodeling, long range chromatin interactions, and gene silencing^7–12^. Here, we use single-molecule methods to provide a biophysical explanation for such findings.

### The SMCHD1 homodimer is flexible and dynamic

We show for the first time, that apart from the closed double rod conformation observed by a previous negative stain electron microscopy study of mouse SMCHD1^19^, human SMCHD1 is able to adopt a variety of conformations. Furthermore, under physiological conditions SMCHD1 is flexible and dynamic. The high flexibility has likely precluded analysis by structural means such as X-ray crystallography and Cryo-EM.

The lengthy linker connecting the hinge and ATPase domains of SMCHD1 is largely uncharacterized and predicted to be **β**-stranded^37^. In our AFM analysis we note that this linker region is flexible, as a SMCHD1^△CC^ truncation mutant involving only the removal of the coiled-coil domain can still adopt various conformations (Fig 5a). Unexpectedly, we note a central density within the linker domain which was observed in all SMCHD1 conformations observed (Supplementary Fig. 1b), the functional characteristics of which remain unclear. It is also apparent that the linker domain can vary its length and affect the overall length of SMCHD1 (Supplementary Fig. 1c, d). Interestingly, AlphaFold2 predicts the SMCHD1 homodimer to be round-shaped rather than rod-shaped, with the ATPase (N-terminus) and hinge (C-terminus) domains in close spatial proximity (Supplementary Fig. 6)^38,39^. A previous study of condensin similarly showed that the hinge domain and ABC-type ATPase head domains can come into close proximity due to the flexibility of the coiled-coil linker^27^. This is somewhat similar to what we observe with the P-shape conformation of SMCHD1, in which we observe interactions in the folded arm between the ATPase domain either with the hinge domain or the opposite arm. Currently the physiological functions of these interactions are unclear, but as the compaction rate in magnetic tweezers is lower for SMCHD1^△link^, we suggest that the flexibility conferred by the linker domain is related to a function of SMCHD1 in compacting DNA and forming DNA clusters. The possible physical interaction between the hinge and ATPase domains may also play a role by both domains independently binding different DNA molecules and bringing them into close proximity for compaction.

Furthermore, in contrast to the coiled-coil domains which make up the linker of canonical SMC core proteins, the ∼22 kDa of coiled-coil regions adjacent to the C-terminus hinge domain of SMCHD1 are relatively minor and we show that their removal does not disrupt the flexibility of the SMCHD1 homodimer (Fig. 5a). Instead, we show that the coiled-coils are important for interaction with LRIF1 (Fig. 5d). LRIF1 also possesses a coiled-coil domain necessary for interaction with SMCHD1^31^. This suggests that the interaction is likely through coiled-coil domains of both proteins. As mutations in both SMCHD1 and LRIF1 have been implicated in FSHD^5,32^, our study provides further insights into their physical interaction.

### ATP regulates the compaction of DNA and reconstituted nucleosomes by SMCHD1

Via both AFM and magnetic tweezers, we show that full-length SMCHD1 compacts DNA and creates protein-DNA clusters without ATP. We used multiple sample deposition methods and various ionic conditions to demonstrate unambiguously that the formation of dense SMCHD1-DNA clusters is a reproducible and physiologically relevant phenomenon (Fig. 2). While SMCHD1 is classified as a non-canonical SMC protein, the DNA compaction phenomenon we observe is reminiscent of the canonical SMC complexes; cohesin, condensin and SMC5/6 have been shown to extrude DNA loops in an ATP-dependent manner^13–15^, although ATP-independent compaction of DNA has also been observed under certain conditions for SMC proteins^40–42^. Canonical SMC proteins can also compact DNA stretched under low forces, as shown by cohesin (0.6 pN and below)^14^, condensin (0.75 pN)^43^, and SMC5/6 (∼0.5 pN)^15,29^. Similarly, SMCHD1 showed the most robust compaction at stretch forces of 0.5 pN and below, with steps in the compaction traces. Given that SMCHD1 does not require ATP for DNA compaction, we suggest that the steps are induced via bridging adjacent DNA segments rather than loop extrusion (Fig. 3).

Perhaps the most significant difference in SMCHD1 compared to the canonical SMC proteins is the presence of a GHKL ATPase domain instead of an ABC-type ATPase domain. The GHKL ATPase of SMCHD1 has a k_cat_ of 0.6 min^-1^ ^44^, while ABC-type ATPases in SMC complexes have a k_cat_ closer to 60 min^-1^ ^13,45^, a difference of 100-fold. ATP binding and hydrolysis are key steps in loop extrusion by SMC complexes^26,46^. As low turnover enzymes, GHKL ATPases are unlikely to generate enough energy for loop extrusion. GHKL ATPase domains are present in a myriad of proteins with various functions, of which the most relevant to the present study is *C. elegans* MORC1 which can also compact DNA^16^.

Interestingly, we find an ATP-dependent effect in both AFM and magnetic tweezers. In magnetic tweezers, the compaction rate in the presence of ATP is reduced (Fig. 6f, g), whilst in AFM using both naked DNA and reconstituted nucleosomes, clusters formed in the presence of ATP are visibly reduced in volume and more disperse compared to the ATP-null condition (Fig 7d, 8c); this phenomenon is also validated with the use of a polynucleosome association assay which demonstrates a decreased degree of nucleosome sedimentation in the presence of ATP (Fig. 7e). We speculate that ATP binding plays a regulatory role in the control of cluster sizes, preventing clusters from getting too large. ATP binding or hydrolysis could potentially cause SMCHD1 to disengage from DNA/chromatin targets. However the exact details for such a model remains to be elucidated. Given the roles of the ATPase domain of SMCHD1 in localizing to genomic targets such as to the inactive X^19,47^, we also do not preclude a role of SMCHD1 ATPase activity in genomic localization.

### SMCHD1 compacts both naked DNA and reconstituted nucleosome arrays

We observe that SMCHD1 can compact both naked DNA and nucleosome arrays (Fig. 2, 7, 8). Whether or not this phenomenon involves direct interaction between SMCHD1 and histones remains to be uncovered, as SMCHD1 may exert its compaction through binding to the DNA within nucleosome arrays. It is unclear if SMCHD1 directly binds histones and/or histone modifications; Peptide pull-downs using H3K9me3 peptides do not pull down purified full-length mouse SMCHD1, but are able to pull down SMCHD1 from nuclear extracts^19^, and this may be attributed to the H3K9me3-Hp1γ-LRIF1-SMCHD1 interaction axis, which allows SMCHD1 to bind H3K9me3 chromatin sites^19^. Therefore, while we observed a direct compaction phenomenon for SMCHD1 and DNA/nucleosome arrays, it is also important to consider histone modifications and/or other direct or indirect binding partners such as Hp1γ-LRIF1, or the multitude of genomic factors involved in X-chromosome inactivation, that may regulate SMCHD1’s compaction activity.

### DNA compaction as a mechanism for epigenetic regulation and chromatin interactions

Our finding that SMCHD1 compacts DNA provides insights towards its roles as an epigenetic regulator. DNA compaction could have a direct role towards altering chromatin landscapes. For instance, SMCHD1 has been attributed to DNA repair although its precise mechanism remains unclear. SMCHD1 is involved in telomeric DNA damage signalling possibly via altering telomere structure to promote the recruitment of downstream DNA repair proteins^48^. It is thus plausible that such structure alteration is directly performed by SMCHD1. SMCHD1 has also be shown to directly bind and maintain a repressed chromatin structure at the subtelomeric D4Z4 macrosatellite repeats to prevent expression of the DUX4 protein which confers muscle toxicity, with mutations in SMCHD1 attributed to the FSHD2 disorder^5,49^. Such a repressed chromatin structure could be a direct result of the DNA compaction activity of SMCHD1.

The DNA compaction activity of SMCHD1 provides a mechanistic insight towards chromatin interactions and merging of chromatin compartments. On the inactive X chromosome, SMCHD1 acts as a bridge for chromatin regions marked with H3K27me3 and H3K9me3^31^, and has been shown to merge chromatin compartments on the inactive X and mediate long-range interactions that traverse topological associated domain (TAD) boundaries^8,9,11^. Our observations that SMCHD1 compacts DNA by bridging multiple adjacent DNA segments provides a potential mechanism for how it may facilitate chromatin mixing. SMCHD1 has also been proposed to act as an insulator by blocking access of proteins such as CTCF or other epigenetic regulators such as TET proteins^23,50^. Our observation of dense clusters formed by SMCHD1 and DNA supports the idea that SMCHD1-bound DNA forms a domain that excludes the entry of other proteins.

Our observation that SMCHD1 forms dense clusters with a significant central intensity suggests a biophysical mechanism (Fig. 9) akin to what has been described as polymer-polymer phase separation or bridging-induced phase separation^51^. SMCHD1 can bind single DNA molecules, and the dense clusters formed suggests that SMCHD1 proteins bridge multiple DNA molecules. Such a phenomenon is possible only when the protein possesses multiple points of contact with DNA, which in this study we present as the hinge and ATPase domain, although it is also possible that other regions of SMCHD1, such as the uncharacterized linker region, also possesses DNA binding activity. As SMCHD1 progressively bridges more DNA molecules, it causes more SMCHD1 particles to bind. This saturates the available DNA binding sites within the cluster and thus gives the appearance of a highly dense cluster (Fig. 9). Polymer-polymer phase separation has been used to describe the behaviour of cohesin in the absence of ATP and it remains to be seen if other chromatin-associated proteins also use this mechanism^40^.

**Fig. 9.**
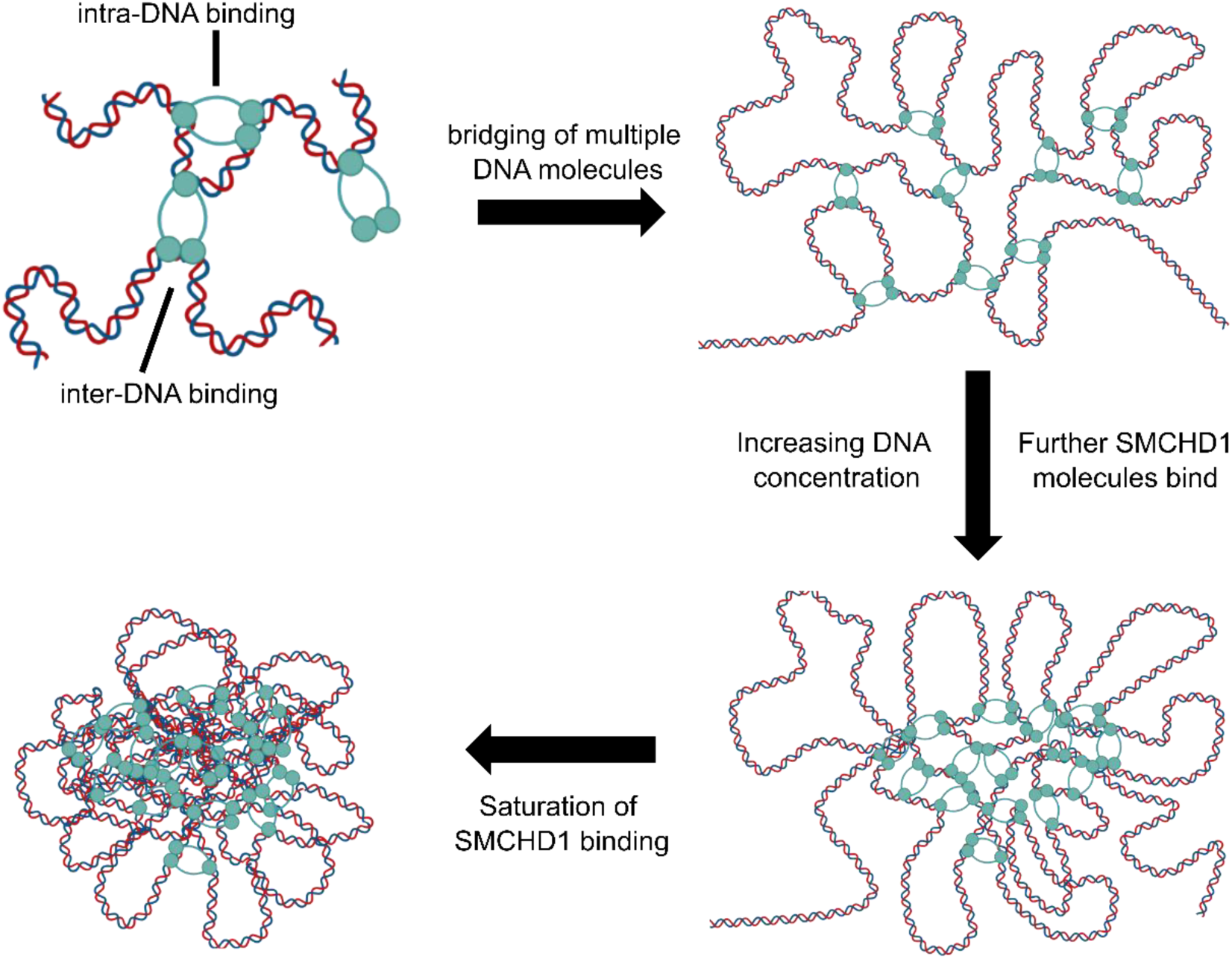
Model for SMCHD1-induced DNA clusters. The progressive binding and bridging of multiple DNA molecules by SMCHD1 increases the local concentration of DNA within the cluster that attracts further SMCHD1 molecules to bind. Given sufficient concentration, DNA binding sites within the cluster eventually become saturated with SMCHD1 binding, giving the appearance of the highly dense cluster.

In summary, we have provided the first biophysical insights towards the epigenetic mechanisms of SMCHD1 by showing it creates dense clusters through compacting DNA. A number of open questions still remain, for example how SMCHD1 is specifically recruited to its genomic targets. It is possible that the ATPase domain is involved in such a function, given its requirement for chromatin localization^19,47^; the precise biochemical mechanism of such a function however, remains to be studied. Secondly, it is unclear if the compaction activity of SMCHD1 could be altered by DNA sequence specificity or by chromatin marks such as histone modifications. Lastly, it is also crucial to understand if FSHD and BAMS mutations would abrogate the *in vitro* DNA or chromatin compaction activity of SMCHD1. Such pursuits would be promising avenues towards developing therapies for these disorders.

## Acknowledgements

We thank all members of the Xue lab for constructive suggestions and thoughtful critiques, and Sho Goh for critical reading of the manuscript. This work was supported by a PYP start-up grant (SX) and the Ministry of Education, Singapore (T2EP30122-0015 to SX). JN was a graduate scholar in receipt of a research scholarship from the National University of Singapore. The authors thank Structural Biology Lab 5 of the Department of Biological Sciences, National University of Singapore, for the kind provision of protein purification facilities.

## Author Contributions

JN, JY and SX designed the research. JN conducted the experiments. JN, XZ, XL, AS, LN, JY and SX analyzed the data. SX supervised the study. JN, JY and SX wrote the paper with inputs from all authors.

## Data availability

Data are available within the article or source data file or from the corresponding author upon reasonable request.

## Code availability

MATLAB code to analyse step sizes in magnetic tweezers has been previously published and is accessible (https://github.com/jacobkers/Loeff-Kerssemakers-et-al-AutoStepFinder).

## MATERIALS AND METHODS

### Cloning and construction of DNA substrates

Full-length SMCHD1 and its mutants, the ATPase domain (aa25-581 or aa25-702) and hinge domain (aa1682-1898) were cloned into a pFastBac vector (Invitrogen) and added with a N-terminus polyhistidine tag with TEV cleavage site for subsequent affinity chromatography. Positive clones were confirmed by Sanger sequencing and transformed into DH10Bac for Bacmid construction.

For AFM DNA substrates, pUC19 DNA was digested at a single site and purified via gel extraction and PCR clean up (Qiagen). The 48.5 kb lambda DNA (NEB) was used as a DNA substrate for AFM without any prior modification. The biotin digoxigenin-labelled DNA substrate for magnetic tweezers experiments was constructed based on the previously published megaprimer approach^52^. Briefly, ∼400 bp biotin/digoxigenin-labelled primers were constructed via a PCR reaction with either biotin-16-dUTP or digoxigenin-11-dUTP (Roche). The 400 bp products were gel extracted (Qiagen) and further used as primers to amplify a 7.5kb region of lambda DNA (NEB), with the final resultant 7.5 kb lambda DNA substrate labelled with biotin and digoxigenin at either end. The substrate was gel extracted and further purified with PCR clean-up (Qiagen).

### Purification of SMCHD1 constructs

Production of baculovirus was conducted according to Bac-to-Bac baculovirus expression system protocols (Thermo Fisher). Bacmid DNA was transfected into SF9 cells with Cellfectin reagent (Gibco) to obtain P0 baculovirus. Baculovirus was then amplified successively to obtain P1 and P2 stocks. For protein expression, 500 ml of SF9 cell culture (2 million cells/ml) was infected with 5 ml of P2 baculovirus and incubated with shaking for 72 hours at 27 ℃ and 120 RPM. Cells were harvested with centrifugation and subsequently flash frozen and stored in -80 ℃.

Purification of full-length SMCHD1 and its truncation mutants (SMCHD1^△CC^, SMCHD1^△link^) was conducted as follows. Cell pellets were resuspended in lysis buffer (20 mM Tris pH 8.0, 150 mM NaCl, 10% Glycerol, 0.2% NP-40, 10 mM imidazole, 0.5 mM TCEP) containing 10U/ml benzonase nuclease (Sigma) and 2 mM MgCl_2_ to digest nucleic acids. The lysate was cleared by two successive rounds of centrifugation at 20,000 RCF and 4 ℃ for 1 hour each, and loaded onto an affinity column containing Ni-NTA agarose (Qiagen), and subsequently incubated with rocking for 1 hour at 4 ℃. Beads were washed three times with wash buffer (20 mM Tris pH 8.0, 500 mM NaCl, 10% Glycerol, 20 mM imidazole, 0.5 mM TCEP) and eluted in buffer containing 250 mM imidazole. The elute was dialyzed into FPLC buffer (20 mM Tris pH 8.0, 150 mM NaCl, 10% Glycerol, 0.5 mM TCEP) and added with TEV protease and 0.5 mM EDTA for overnight cleavage of the 6xhis tag. The elute was then run on a Superose 6 increase column (Cytiva) in FPLC buffer; peak fractions were pooled, flash frozen and stored in -80 ℃. The individual SMCHD1 hinge and ATPase domains were purified similarly, with the exception of a use of a Superdex 75 increase column (Cytiva) for size exclusion chromatography. Purity of proteins was assessed with 10% SDS-PAGE; purity of full-length SMCHD1 was further assessed with Western Blot using anti-SMCHD1 (Atlas Antibodies HPA039441).

### Sample preparation for atomic force microscopy

For AFM sample preparation involving magnesium as counterion (Fig. 2a-c, Fig. 3, Fig. 4, Fig 5b, e), SMCHD1 and DNA substrates were incubated for 10 mins in binding buffer (20 mM Tris-Cl pH 8.0, 50 mM NaCl, 2.5 mM MgCl_2_, 0.5 mM TCEP) at room temperature. The mixture was then diluted five-fold with deposition buffer (20 mM Tris-Cl pH 8.0, 10 mM MgCl_2_, 0.5 mM TCEP) and immediately deposited on freshly cleaved mica. Following an incubation time of two minutes, the sample was washed gently with 3 ml of sterile-filtered MilliQ water and dried with a gentle stream of nitrogen gas. For AFM sample preparation without counterions (Supplementary Fig. 2b), SMCHD1 and DNA substrates were incubated for 10 mins in binding buffer (20 mM Tris-Cl pH 8.0, 50/150 mM NaCl, 0.5 mM TCEP) at room temperature, before being deposited on freshly cleaved mica without any prior modification. For AFM sample preparation involving APTES and glutaraldehyde (Fig. 2d-g, Supplementary Fig. 2a) as previously described^22^, freshly cleaved mica was incubated with 0.1% APTES for 15 minutes and rinsed with MilliQ water, then incubated with 1% glutaraldehyde for 15 mins and rinsed with MilliQ water. SMCHD1-DNA incubations were then deposited on this treated mica similarly as AFM sample preparation without counterions.

For sample preparation of liquid AFM imaging of individual SMCHD1 particles, SMCHD1 was deposited on freshly cleaved mica in imaging buffer (20 mM Tris-Cl 8, 150 mM NaCl, 0.5 mM TCEP, 10 mM MgCl_2_). The sample was incubated for 10 seconds and gently washed with 3 ml of imaging buffer. It was then placed without drying into the AFM imaging chamber. All buffers for AFM experiments were prepared daily from 1M stock components stored in 4 ℃. For SMCHD1-nucleosome experiments (Fig. 7 and Fig. 8), we used reconstituted 15-197-601 or 20-157-601 nucleosome arrays prepared as described in prior studies^53,54^. Mica was pre-treated with 0.001% poly-L-lysine treated for 15 minutes and rinsed thoroughly with MilliQ water. Reconstituted 15-197-601 or 20-157-601 nucleosome arrays at a final concentration of 1ng/μl were incubated with purified full-length SMCHD1 of indicated concentrations in binding buffer (20 mM Tris-Cl pH 8.0, 50 mM KCl, 0.5 mM MgCl_2_, 1 mM DTT) with or without 2 mM ATP or GTP for 30 minutes at room temperature. The sample was then deposited onto the poly-L-lysine treated mica and left to incubate for 15 minutes. The mica was then washed with 1 ml of ultra-pure water and gently dried with a stream of nitrogen gas. Array-only samples were prepared similarly in the absence of SMCHD1; SMCHD1-only samples were prepared similarly in the absence of nucleosome arrays. For incubations involving the addition of the SMCHD1 inhibitor radicicol (Fig. 8b), 80 nM SMCHD1 was first incubated for one hour with 100 μM final concentration of radicicol prior to incubation with nucleosomes with or without ATP. Sample preparation for volume assessment of individual SMCHD1 particles with or without ATP, in relation to nucleosome experiments (Supplementary Fig. 5d) were conducted in similar buffer conditions using as per nucleosome experiments, using 20 nM SMCHD1. Volume of individual particles were assessed from 2 by 2 µm images.

### Atomic force microscopy

AFM imaging was conducted with a Bruker Dimension FastScan with Nanoscope software and ScanAsyst imaging mode. For dry AFM, FastScan C probes (specifications: 250 kHz resonant frequency, 1.5 N/m force constant) were used; images were typically taken at 2 by 2 µm, 4 by 4 µm or 10 by 10 µm scan areas as indicated, resolution of 512 x 512 pixels, at a scanning speed of 4 Hz per line. For each independent experiment, images were obtained from the same mica sample at different scan locations. Liquid AFM was conducted similarly except with the use of FastScan-D probes. For liquid AFM of individual SMCHD1 particles, we used a scan size of 200 x 200 nm, 256 x 256 pixels, at a scanning speed of 15 Hz per line. AFM images were analysed and processed with Gwyddion software version 2.62.

#### AFM image processing

AFM images were processed with Gwyddion version 2.62 as follows. Raw AFM images were base flattened and data levelled using mean plane subtraction. Polynomial background was then removed and rows aligned using a 1^st^ degree polynomial. Scan line artefacts were removed by correcting horizontal scars. For volume measurements and comparisons (Fig. 2, 5, 8), AFM images were masked using height threshold and raw grain values were exported and plotted. Volume of individual nucleosome clusters (Supplementary Fig. 5) was assessed manually similarly using height threshold. Height thresholds used for volume comparisons are indicated in figure captions. Section profiles of individual SMCHD1 particles (Figure 1c, Supplementary Fig. 1c) were illustrated with Nanoscope analysis software.

### Polynucleosome array association assay

We used a modified polynucleosome self-association assay (Fig. 7e) recently used in a study to assess protein-induced chromatin condensation^36^. Reconstituted nucleosome arrays at a final concentration of 10ng/μl were incubated with purified full-length SMCHD1 of indicated concentrations in binding buffer (20 mM Tris-Cl pH 8.0, 50 mM KCl, 0.5 mM MgCl_2_, 1 mM DTT) with or without 2 mM ATP in a total assay volume of 20 ul. The reaction was incubated at room temperature for 30 minutes before subsequent centrifugation for 10 minutes at 20,000 RCF at room temperature. 10 μl of the resultant supernatant was carefully pipetted out and mixed with 10 μl of SDS-loading buffer (20 mM Tris-Cl pH 8.0, 25% glycerol, 0.25% SDS, 80 mM EDTA, proteinase K (0.5 mg/ml, Thermo Fischer Scientific)). The released DNA fragments were then assessed by 0.8% agarose gel electrophoresis and ethidium bromide staining for visualization. Reactions involving nucleosome array alone were conducted similarly without the addition of SMCHD1. Image analysis of DNA band intensities was conducted with imageJ software. For each independent experiment, DNA intensities as percentages were normalized to the band with the highest intensity.

### Single-molecule magnetic tweezers

For magnetic tweezers flow chamber construction, APTES and glutaraldehyde-treated glass slides were sandwiched to a cover slip with Grace adhesive seals (Sigma) to create channels. The channels were incubated with 20ng/µl anti-digoxigenin for 4 hours at room temperature before extensive washing with 1x PBS. Finally, the channels were passivated in 1x PBS containing 2% BSA and 5 mM BME, and stored in 4 ℃. Prior to experiments, channels were washed extensively with 1x PBS and incubated with 0.01ng/µl of 7.5 kb biotin-digoxigenin DNA substrate for 30 minutes. Channels were washed with 1x PBS to remove free DNA, and added with Dynabeads MyOne Streptavidin T1 beads (Thermo Fisher) for a 20 minutes incubation. Finally, the flow channel was washed with compaction buffer (20 mM Tris-CI pH 8.0, 50 mM NaCl, 0.5 mM TCEP) to remove unbound streptavidin beads. All magnetic tweezers experiments were conducted in compaction buffer; 2 mM MgCl_2_ was included in compaction buffer for experiments involving ATP. For each experimental run, DNA was first stretched at a high force (8 pN) prior to injection of 10 nM SMCHD1. After injection of 10 nM SMCHD1 followed by an incubation period of 10 minutes, the force was lowered to observe compaction. At the end of the run, high force of 8 pN was used to stretch DNA back to observed force-induced decompaction. A home built vertical magnetic tweezers set-up was controlled using LabVIEW program (National Instruments). Extension changes in DNA were monitored via the diffraction pattern of the magnetic bead tracked with a CCD camera capturing ∼160 frames per second. DNA compaction rates were determined by calculating the difference in DNA extension length between the 5% and 95% time points of lowered force of 90 seconds (Fig. 6f), meaning compaction rate was assessed over precisely 81 seconds. DNA extension lengths at the end of the compaction traces were similarly assessed at the end of this 81s timepoint (Fig. 6g). All buffers for magnetic tweezers experiments were prepared daily from 1M stock components stored in 4 ℃. Step-fitting of traces was conducted with the Autostepfinder program under default parameters^33^.

### Statistical Analysis

Statistical tests were conducted with OriginPro 2021b. For magnetic tweezers experiments (Fig. 6, Supplementary Fig. 4), one-way ANOVA with Dunnett’s test for multiple comparisons was used to compare DNA compaction rate (Fig. 6f) and unpaired two tailed t-test for comparing extension lengths (Fig. 6g). Statistics for volume comparisons for AFM experiments were conducted with two tailed t-test. Difference between means was considered significant when p<0.05. Exact p-values are indicated in source data file. For boxplots presented in figures, the middle quartile marks the median, boxes extend to the quartiles, and the whiskers depict the range of the data within 1.5 times of the interquartile range of the median. Where applicable, continuous variables were presented as means with 95% confidence interval. Histograms are fitted with normal distribution.

### ATPase assay

ATPase assay was conducted with Innova Biosciences ATPase assay kit according to the manufacturer’s instructions, in similar buffer to SMCHD1 nucleosome AFM experiments (20 mM Tris-Cl pH 8.0, 50 mM KCl, 0.5 mM MgCl_2_, 1 mM DTT). For each experiment, 80 nM SMCHD1 was assayed in ATP + 100 µM radicicol and ATP-only conditions. Absorbance readings were taken at 600 nm wavelength. Decrease in ATPase hydrolysis rates with radicicol was assessed by normalizing to SMCHD1-ATP and buffer alone with ATP. Three independent experiments were conducted.

### EMSA

FAM-labelled double-strand duplex 37bp DNA (5’ 56-FAM/ CCTAGGCTACACCTACTCTTTGTAAGAATTAAGCTTC 3’, IDT) was incubated with purified hinge or purified ATPase domain in binding buffer (20 mM Tris-Cl pH 8.0, 50 mM NaCl, 2 mM MgCl_2_, 10% Glycerol) and incubated for 30 minutes on ice. The samples were run on a 1.5% TAE agarose gel in TAE buffer at 80V for 1.5 hours at 4 ℃. Gels were imaged with a Biorad Chemi-Doc MP with a fluorescein filter.

### Co-immunoprecipitation

For co-immunoprecipitation of overexpressed constructs, FLAG-tagged SMCHD1 and HA-tagged LRIF1 were transfected into 293T cells using Lipofectamine 2000 (Thermo Fisher). At 48 hours post-transfection, cells were harvested by first washing in ice-cold PBS and lysed in lysis buffer (50 mM Tris-Cl pH 8.0, 150 mM NaCl, 0.5% Triton X) supplemented with cOmplete protease inhibitor cocktail (Roche). Cells were incubated with rocking for 30 minutes at 4 ℃ and subsequently centrifuged at max speed for 30 minutes at 4 ℃ to remove insoluble debris. After collecting an input sample, anti-flag M2 agarose beads (Sigma) were added to the remainder of the soluble lysate and incubated for two hours with rocking at 4 ℃. Beads were then washed three times with lysis buffer and boiled with 2x SDS-PAGE buffer for subsequent western blot analysis. Antibodies used for western blot were anti-flag (Sigma F1804) and anti-HA (Cell-Signaling C29F4). Three independent experiments were conducted.

### Alphafold

Human SMCHD1 dimer was folded by Alphafold v2 Multimer^38,39^. The template date was set to 2022-03-30. Default parameters were used for all other settings.

### Graphics

Several illustrations depicted in figures were created with Biorender^55^.

---

**Supplementary Figure 1.**
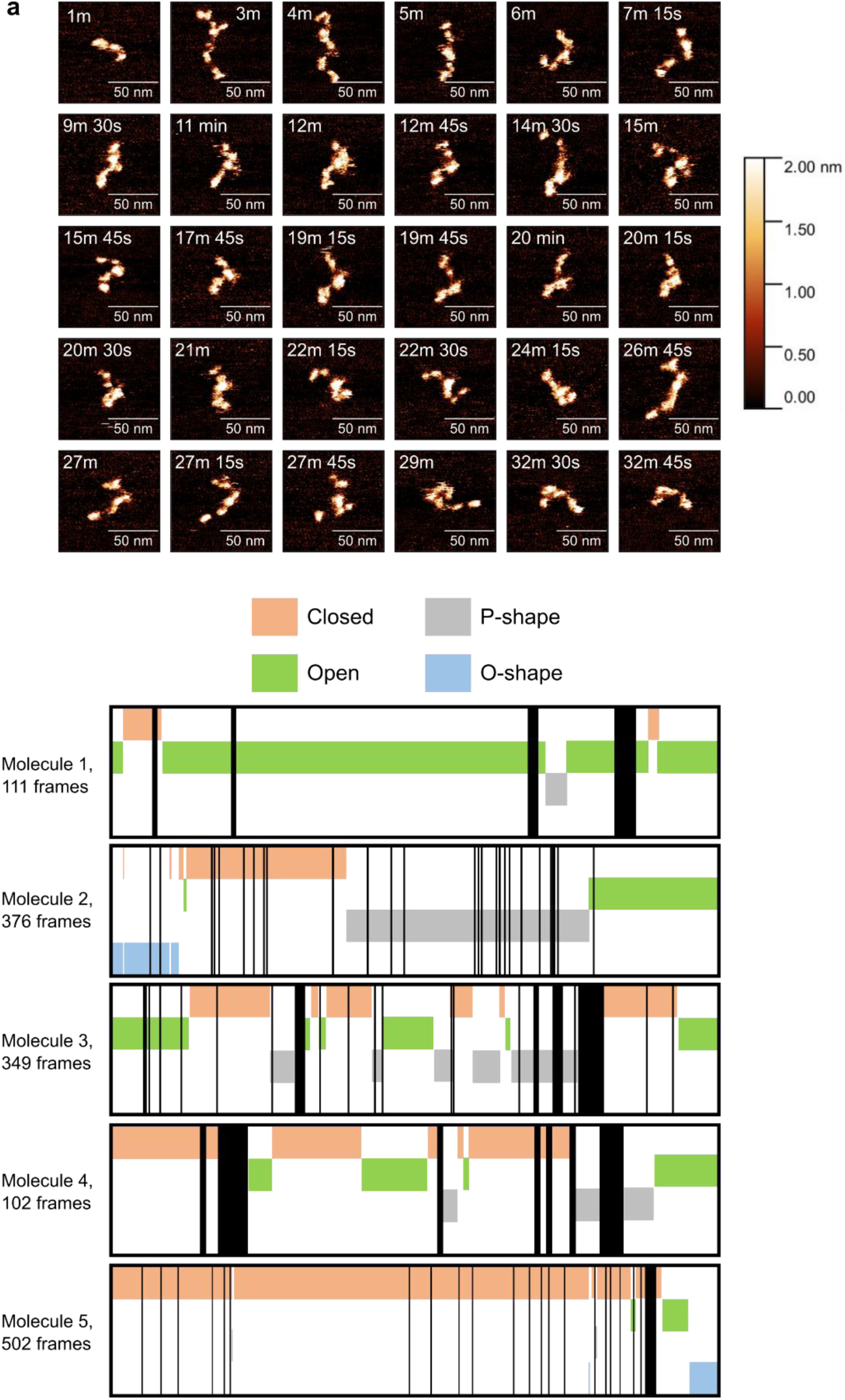
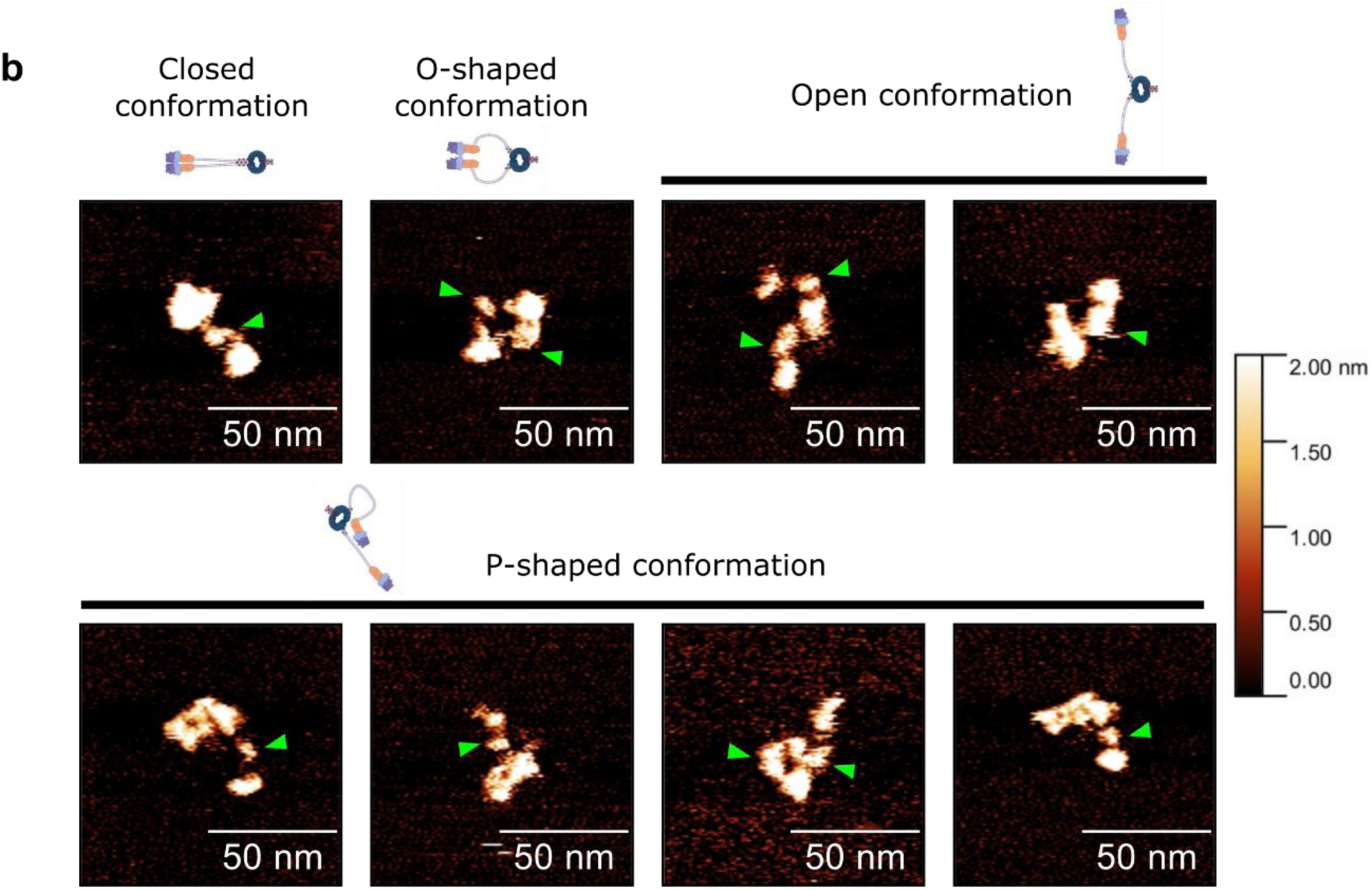
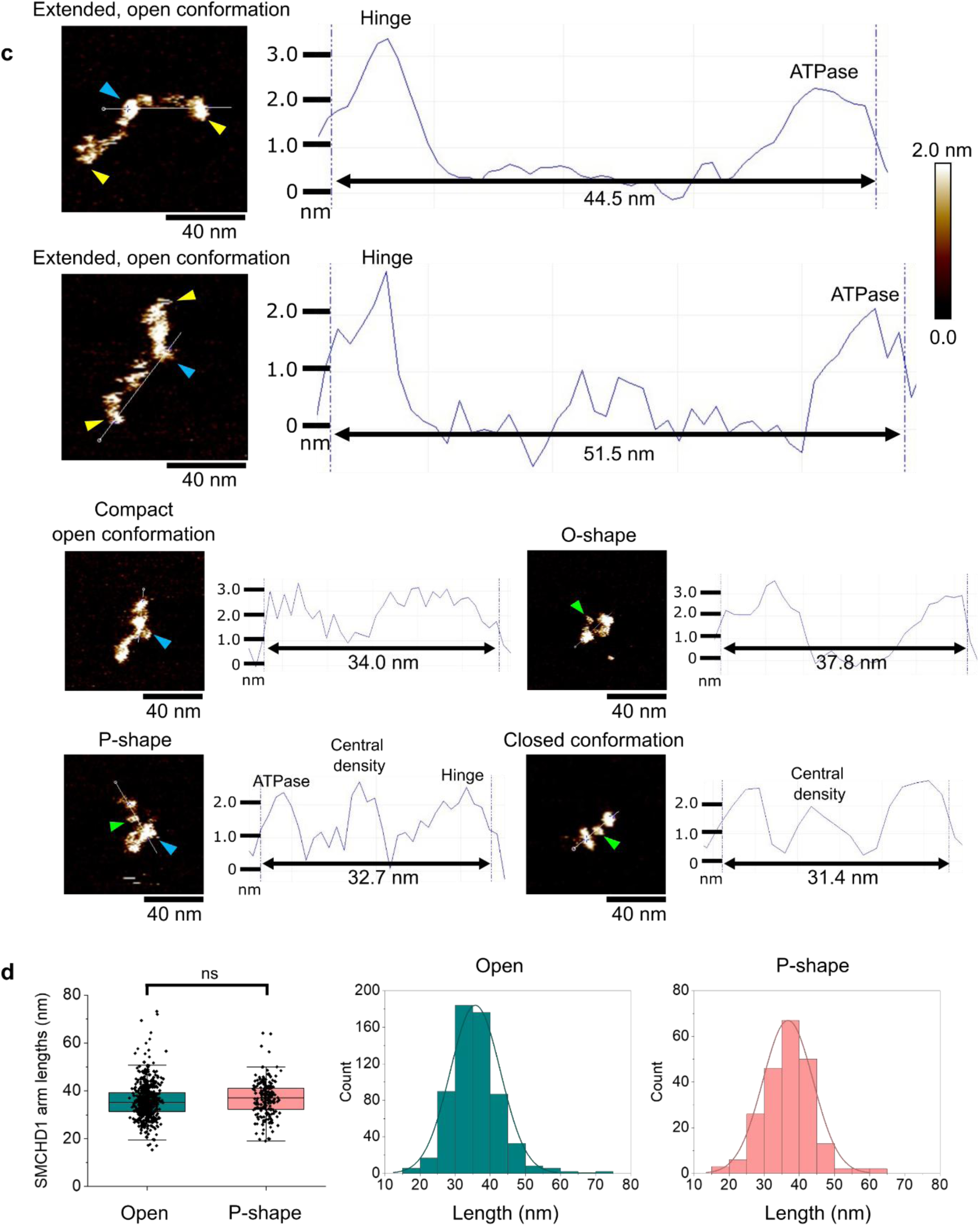
Liquid AFM of SMCHD1. **a**, *Top*, Snapshots of liquid AFM imaging of SMCHD1, related to Figures 1d-1f, and corresponding to molecule 1 in Supplementary Movie 1. All images were taken at an imaging speed of 15 Hz per line. *Bottom*, annotation of conformations for five individual SMCHD1 molecules imaged with liquid AFM. Molecules 1 to 3 correspond to Supplementary Movies 1-3. Black vertical lines indicate frames in which the conformation of SMCHD1 could not be confidently interpreted. See Supplementary Movies 1-3 for the AFM movies of the three molecules. **b**, Further molecules of SMCHD1 in various conformations displaying the central density within the linker region, indicated with a green arrowhead. **c**, Top, section profiles of individual SMCHD1 homodimers to illustrate the varying hinge-to-ATPase distances in different conformations, as mediated by the linker domain. The first two molecules depict the linker domain in extended conformations which correspond to increased end-to-end distance between the hinge and ATPase. Other molecules depict shorter end-to-end distances. Where applicable, blue arrowheads indicate the hinge domain, yellow arrowheads indicate the ATPase domain, while green arrowheads indicate the linker central density. The 2nm z-scale bar depicted applies to all AFM images in this figure. **d**, comparison and distribution of SMCHD1 arm lengths for open and P-shape conformations. For boxplots in **d**, the middle quartile marks the median, boxes extend to the quartiles, and the whiskers depict the range of the data within 1.5 times of the interquartile range of the median; student’s t-test was used for statistical analysis. Histograms were fitted with normal distribution.

**Supplementary Figure 2.**
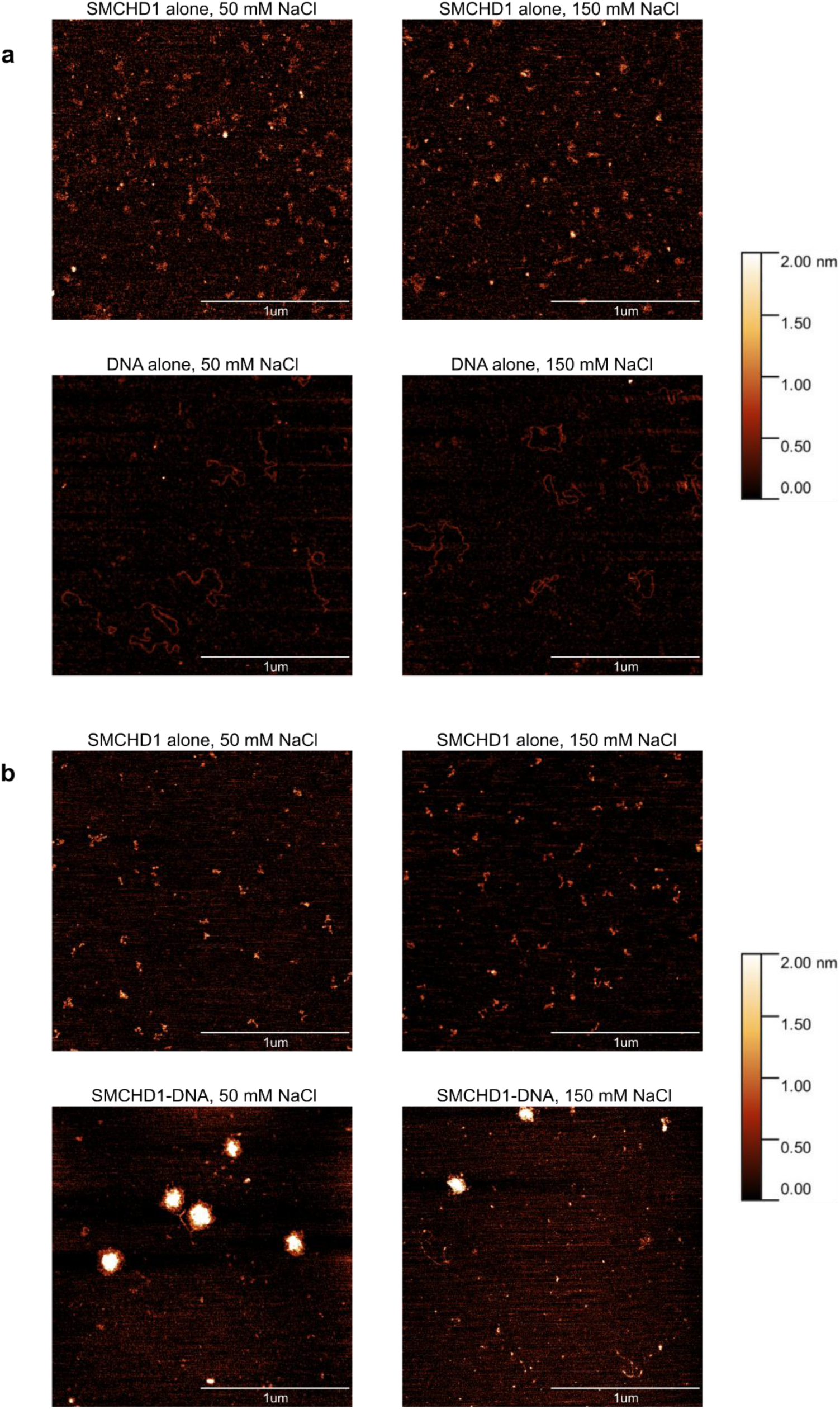
SMCHD1 compacts DNA under various sample deposition conditions. **a**, SMCHD1 alone or DNA alone in buffer containing 50 or 150 mM NaCl, deposited on APTES-glutaraldehyde treated mica, related to Figure 2d. **b**, *Top*, SMCHD1 alone in counterion-free buffer containing either 50 or 150 mM NaCl, deposited on fresh mica. *Bottom*, SMCHD1-DNA clusters in counterion-free buffer containing either 50 or 150 mM NaCl, deposited on fresh mica.

**Supplementary Figure 3.**
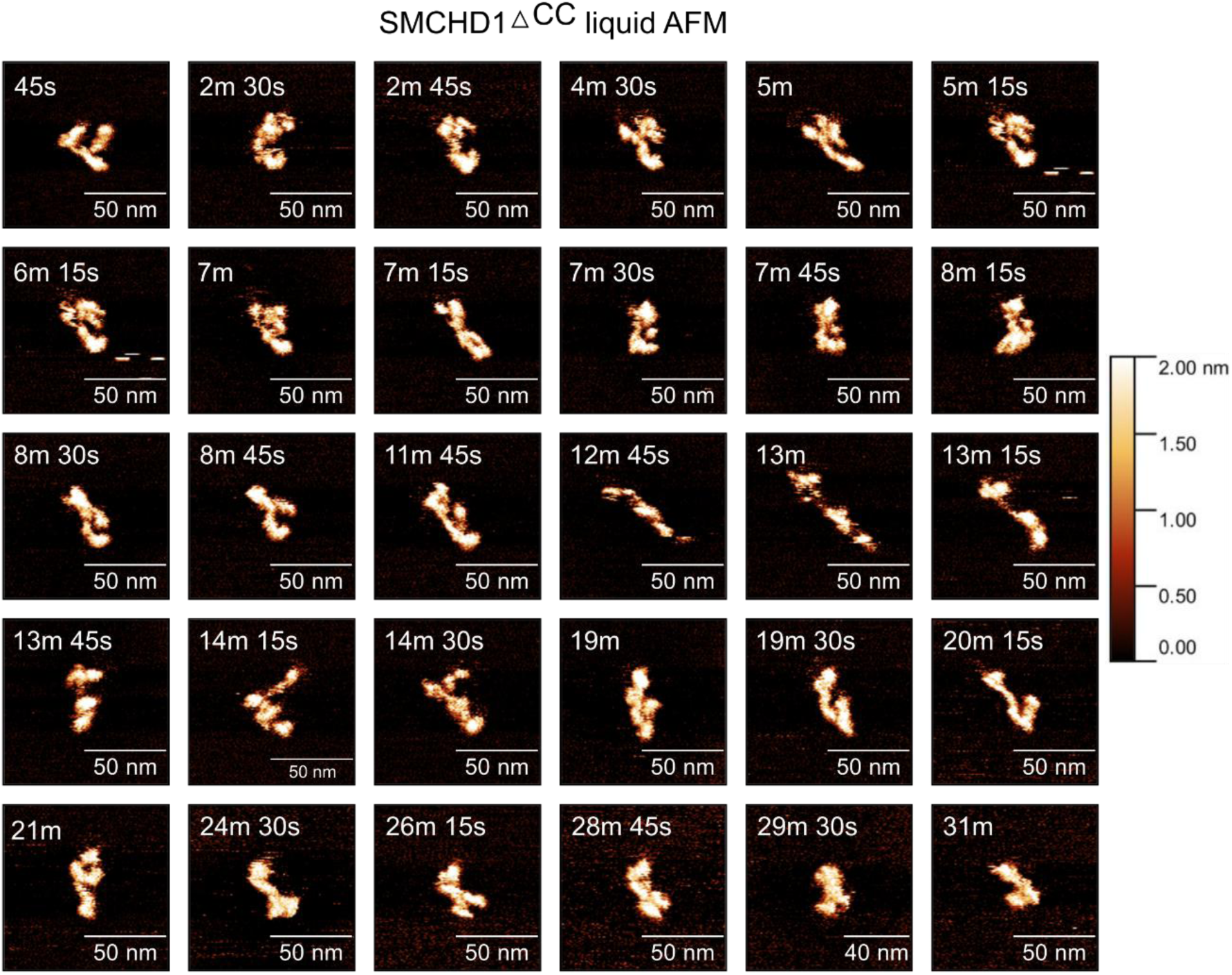
SMCHD1^△CC^ adopts various conformations in liquid AFM. Selected snapshots of liquid AFM imaging of SMCHD1^△CC^ over a period of 30 minutes, related to Fig. 5a. All images were taken at an imaging speed of 15 Hz per line.

**Supplementary Figure 4.**
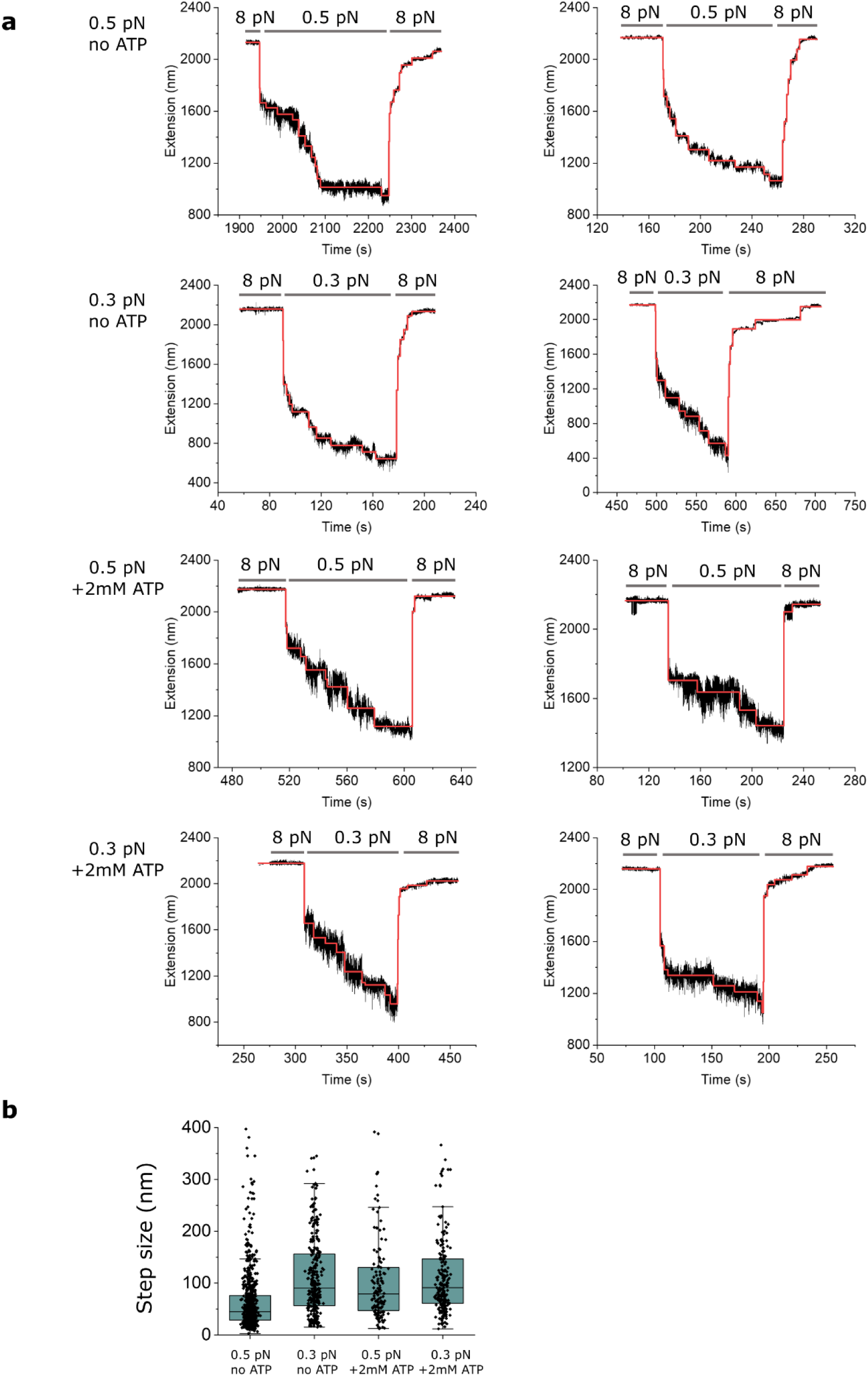
Characterization of step sizes induced by SMCHD1-mediated DNA compaction, related to Figure 6f and 6g. **a**, Representative compaction and decompaction traces of SMCHD1-mediated DNA compaction under 0.5/0.3 pN and in the presence or absence of ATP. Red lines indicate the fit by Autostepfinder^33^. **b**, Boxplot comparison of the compaction step sizes between different compaction conditions (0.5 pN no ATP, n = 704; 0.3 pN no ATP, n = 306; 0.5 pN + 2 mM ATP, n = 134; 0.3 pN + 2 mM ATP, n = 207). The middle quartile marks the median, boxes extend to the quartiles, and the whiskers depict the range of the data within 1.5 times of the interquartile range of the median. Source data is provided as a source data file.

**Supplementary Figure 5.**
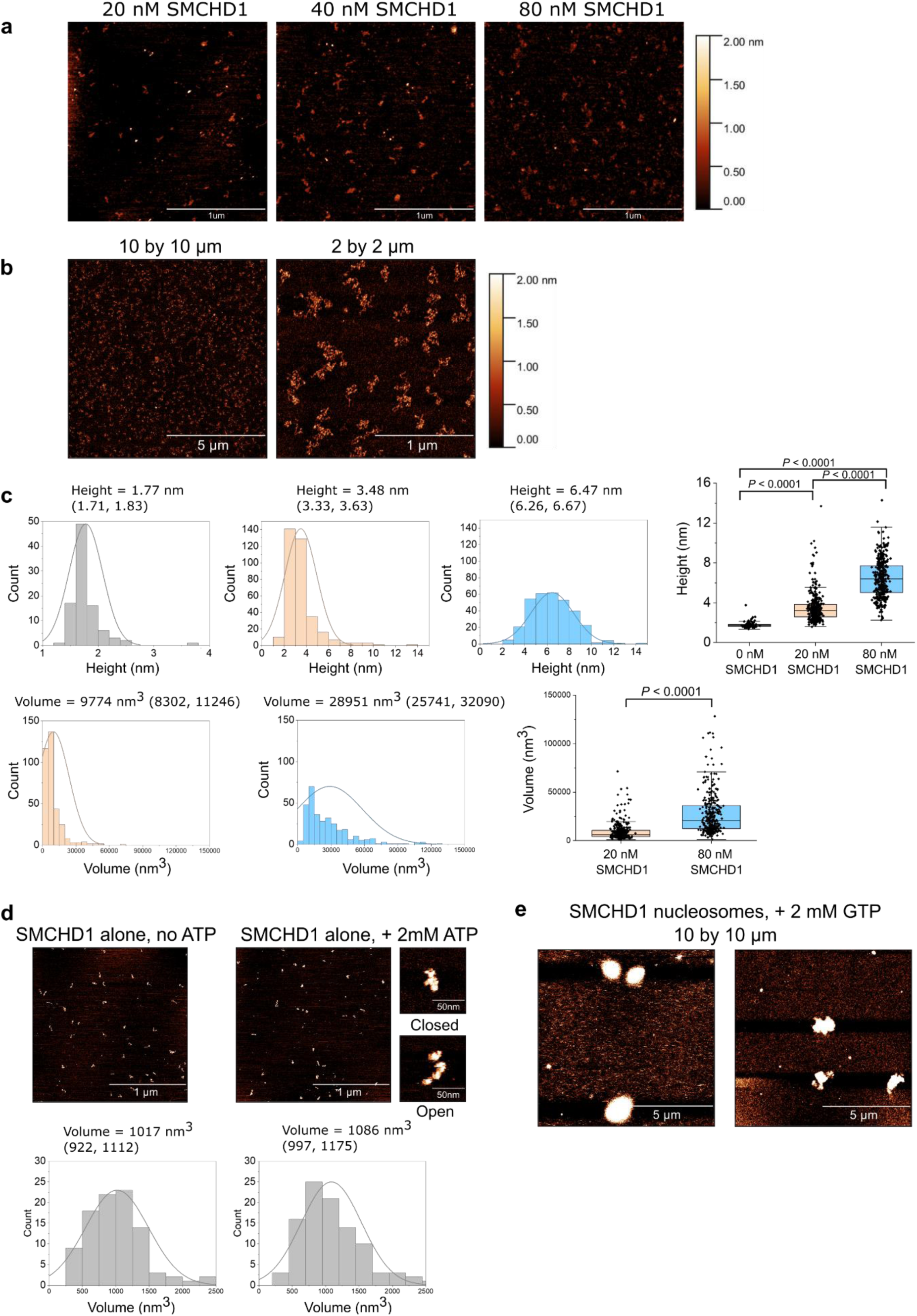
**a**, SMCHD1 alone (20, 40 and 80 nM) deposited on poly-L-lysine treated mica. **b**, 10 by 10 and 2 by 2 µm AFM images of nucleosomes alone in the presence of ATP. **c**, quantification of the height and volume of individual SMCHD1-nucleosome clusters between 20 and 80 nM SMCHD1 in the presence of ATP, n = 95, 349 and 328 for nucleosome alone, 20 nM SMCHD1, 80 nM SMCHD1 respectively. Three independent experiments were conducted for 20 nM and 80 nM SMCHD1. **d**, volume of individual SMCHD1 particles assessed from 20 nM SMCHD1 using liquid AFM, either with or without ATP. Data is presented as means with 95% confidence interval, histogram is fitted with normal distribution. Two representative images of individual SMCHD1 particles in a closed and open conformation are depicted for the ATP condition. **e**, 10 by 10 µm AFM image of 80 nM SMCHD1 nucleosome clusters in the presence of 2 mM GTP. Source data is provided as a source data file.

**Supplementary Figure 6.**
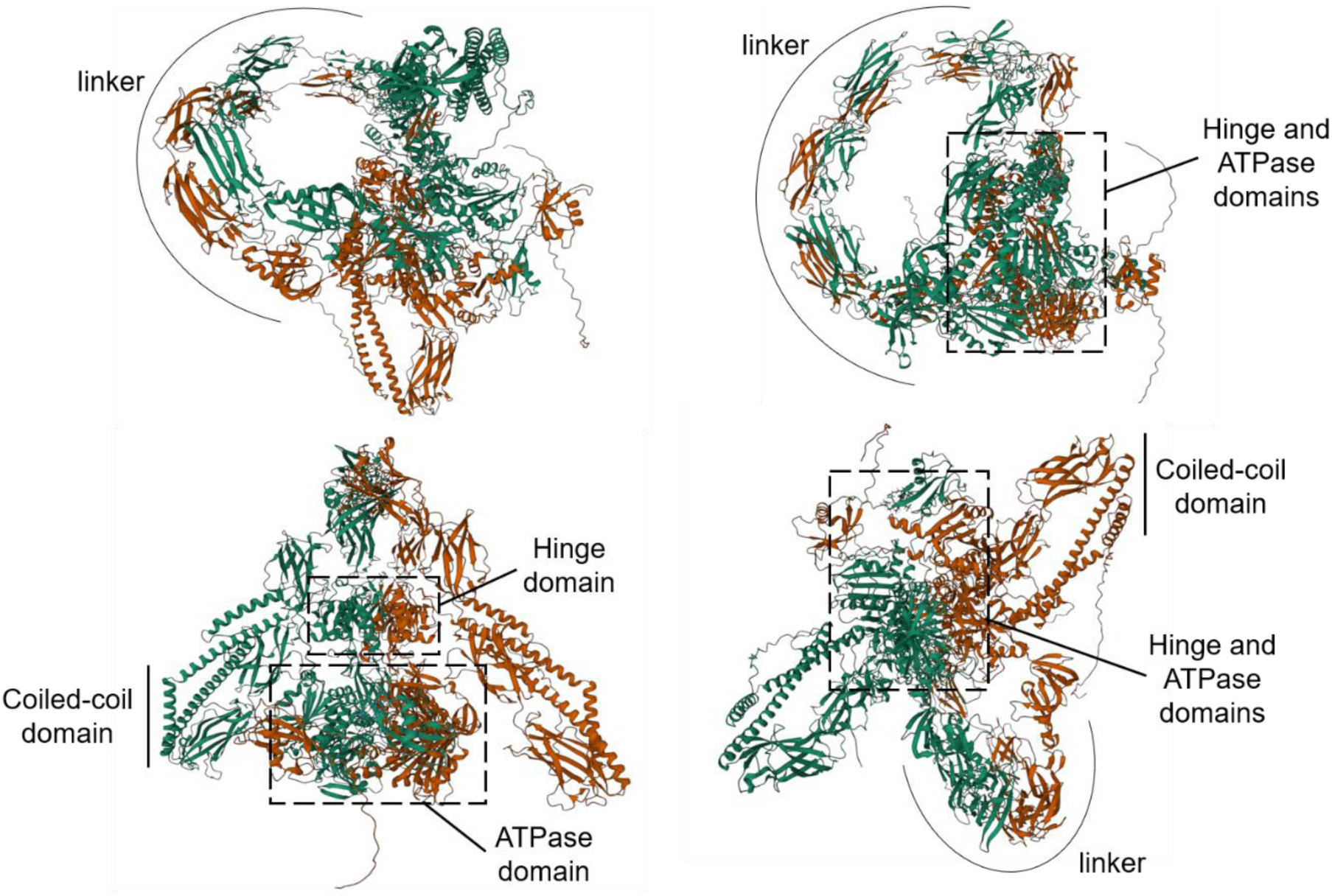
Alphafold prediction of the SMCHD1 homodimer^38,39^. Green and red colours indicate each monomer of the homodimer. From this model, it is clear that the lengthy linker region of SMCHD1 is not rod-like and rigid as previously thought, but rather adopts a circular conformation that brings the Hinge and ATPase domains into close proximity.

